# Organelle membrane-associated proteins recruit cGAS via phase separation to facilitate its membrane localization

**DOI:** 10.1101/2025.08.01.668185

**Authors:** Chengrui Shi, Chaofei Su, Kaixiang Zhang, Hang Yin

**Author notes:** Corresponding author: Hang Yin.

## Abstract

Double-stranded DNA is recognized as a danger signal by cyclic guanosine monophosphate-adenosine monophosphate synthase (cGAS), triggering innate immune responses in mammals. As a DNA sensor, cGAS is generally believed to distribute in the cytoplasm, whereas alternative subcellular localization of cGAS, including cytoplasmic membrane and nucleus, is important to regulate its activity. However, it remains obscure whether cGAS could localized to organelle membrane and the mechanism has yet to be uncovered. Our study reveals that cGAS could localize to the endoplasmic reticulum, Golgi apparatus, and endosomes upon DNA challenge. We identified that the post-translational modification enzymes ZDHHC18 and MARCH8, through their intrinsically disordered regions (IDRs), facilitate the binding of cGAS to the Golgi and endosome, respectively. These IDRs phase separated to recruit cGAS and double-stranded DNA (dsDNA) into biomolecular condensates, suppressing cGAS activity and downstream signaling pathways. These findings highlight the regulatory mechanisms of cGAS activity through the spatial organization, providing new insights into the modulation of innate immune responses.

## Introduction

As a pattern-recognition receptor (PRR), cGAS recognizes double-stranded DNA (dsDNA) and plays essential roles in innate immunity (Sun et al., 2013). Upon DNA binding, cGAS becomes enzymatically active and produces cyclic GMP-AMP (cGAMP), a second messenger which binds stimulator of interferon genes (STING, also known as TMEM173), leads to its conformational changes that trigger oligomerization and activation (Burdette et al., 2011; Gao et al., 2013; Shang et al., 2019; Zhang et al., 2019). After trafficking from the endoplasmic reticulum (ER) to Golgi, activated STING recruits TANK-binding kinase 1 (TBK1) to promote TBK1 autophosphorylation, STING phosphorylation at Ser366 and subsequent recruitment and phosphorylation of interferon regulatory factor 3 (IRF3) (Dobbs et al., 2015; Ishikawa et al., 2009; Saitoh et al., 2009). Phosphorylated dimeric IRF3 then translocate into the nucleus and induces the expression of type I interferons (IFNs) and several other inflammatory cytokines and chemokines (Chen et al., 2016b).

The precise subcellular localization of cGAS is critical for its function, with evidence supporting its presence at the plasma membrane and in the nucleus. These distinct localizations are pivotal for the regulation of cGAS activity, with membrane localization minimizing self-DNA stimulation and nuclear localization inhibiting its activation by interacting with nucleosome (Barnett et al., 2019; Boyer et al., 2020; Cao et al., 2020; Pathare et al., 2020). Despite the well-documented roles of cGAS at the plasma membrane and in the nucleus, its potential localization to other organelles such as the endoplasmic reticulum (ER), Golgi apparatus, and endosomes remains unexplored.

Recent studies have highlighted the importance of IDRs in proteins for facilitating liquid-liquid phase separation (LLPS), which is essential for the formation of biomolecular condensates. These condensates play crucial roles in organizing cellular biochemistry, including the spatial regulation of signaling pathways (Banjade and Rosen, 2014; Chen et al., 2016a; Li et al., 2012; Zeng et al., 2016; Zhao and Zhang, 2020). Binding of dsDNA to cGAS induces LLPS, forming aggregates that enhance the synthesis of 2’3’-cGAMP, which subsequently triggers a STING-dependent immune response (Du and Chen, 2018). The LLPS of cGAS-DNA complexes protects the DNA from degradation by the cytosolic exonuclease TREX1 (Zhou et al., 2021). Proteins such as G3BP1 and USP15 promote cGAS phase separation, while viral protein ORF52 antagonizes it to modulate cGAS activity (Bhowmik et al., 2021; Shi et al., 2022a; Zhao et al., 2022).

In addition to occurring in the cytoplasm or nucleus, phase separation on cell membranes helps regulate the dynamics and efficiency of signal transduction (Huang et al., 2016). Phase separation could also occur on organelle membranes. Argonaute (AGO) proteins undergo lipid-mediated phase separation on the endoplasmic reticulum (ER) membrane, which are crucial for the quality control of nascent peptides (Gao et al., 2022). Besides, calcium transients on the ER surface induce the liquid-liquid phase separation of the protein FIP200, which aggregate to define the sites where autophagosomes are initiated (Zheng et al., 2022). Phase separation also played an essential role in facilitating the transport of messenger ribonucleoprotein particles (mRNPs) and endosomes along microtubules (Baumann et al., 2012). Phase separation is also important for intracellular long-distance transport utilizing lysosomes (Liao et al., 2019). However, it remains unclear whether liquid-liquid phase separation can occur on other organelle membranes, and the regulatory role in cGAS function has yet to be discovered.

Our research reveals that cGAS can localize to the ER, Golgi apparatus, and endosomes. We identified that ZDHHC18 and MARCH8, enzymes that catalyze post-translational modifications of cGAS, utilize their IDRs to anchor cGAS to the Golgi and endosome, respectively. Both ZDHHC18 and MARCH8 undergo phase separation through their IDRs in cells and in vitro, recruiting cGAS and double-stranded DNA (dsDNA) into these biomolecular condensates. This localization and recruitment lead to the suppression of cGAS activity and downstream signaling, highlighting a novel regulatory mechanism of cGAS by spatial sequestration. In summary, our findings provide new insights into the subcellular localization and regulation of cGAS, emphasizing the critical roles of ZDHHC18 and MARCH8 in modulating cGAS activity through phase separation.

## Model

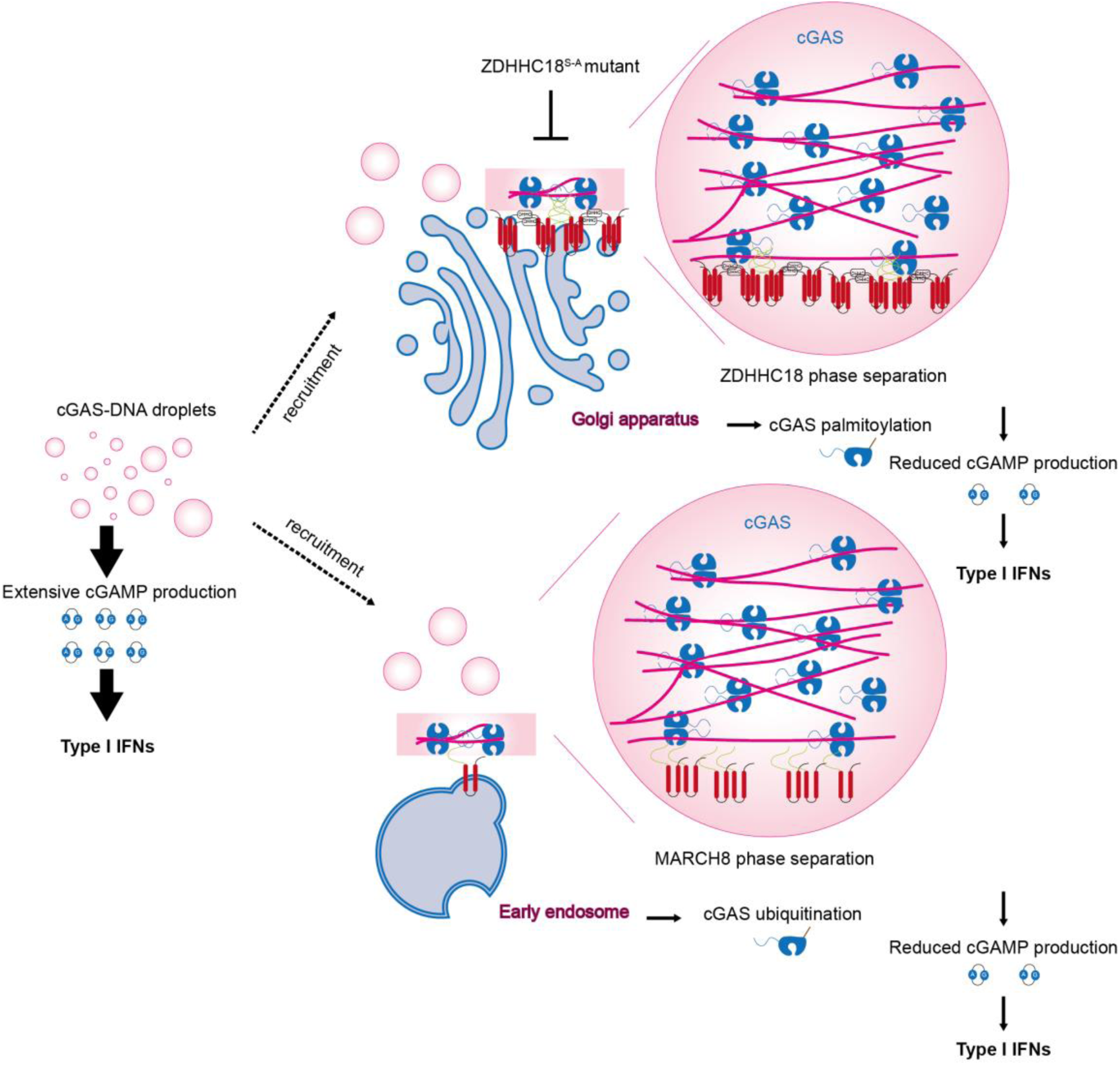

## Results

### cGAS was associated with organelle membranes

To determine the localization of cGAS, we performed subcellular fractionation and utilized Optiprep density gradient centrifugation to isolate fractions containing cGAS without and with Herring testis DNA (HT-DNA). As shown in Fig. 1a, in its steady state, cGAS was broadly distributed and partially overlapped with ER fractions. Upon HT-DNA stimulation, cGAS accumulated in lighter fractions, overlapping with GM130, Calnexin and EEA1, suggesting association with the Golgi, endosome and enhanced association with ER. To exclude nonspecific interactions during cell lysis, we added 1% Triton X-100 to the lysis buffer (Barnett et al., 2019). The results showed that treatment with Triton X-100 led to marked redistribution of ER and Golgi markers: Calnexin shifted to lighter fractions (yellow dashed triangle), and GM130 exhibited a biphasic distribution (green dashed triangle). Correspondingly, in the presence of HT-DNA, cGAS redistributed to these altered organelle fractions. Notably, cGAS remained present in the solubilized ER fractions (yellow solid triangle) and was also retained in the redistributed Golgi fractions (green solid triangle), indicating its continued interaction with membrane-associated compartments despite detergent-mediated reorganization.

**Fig. 1.**
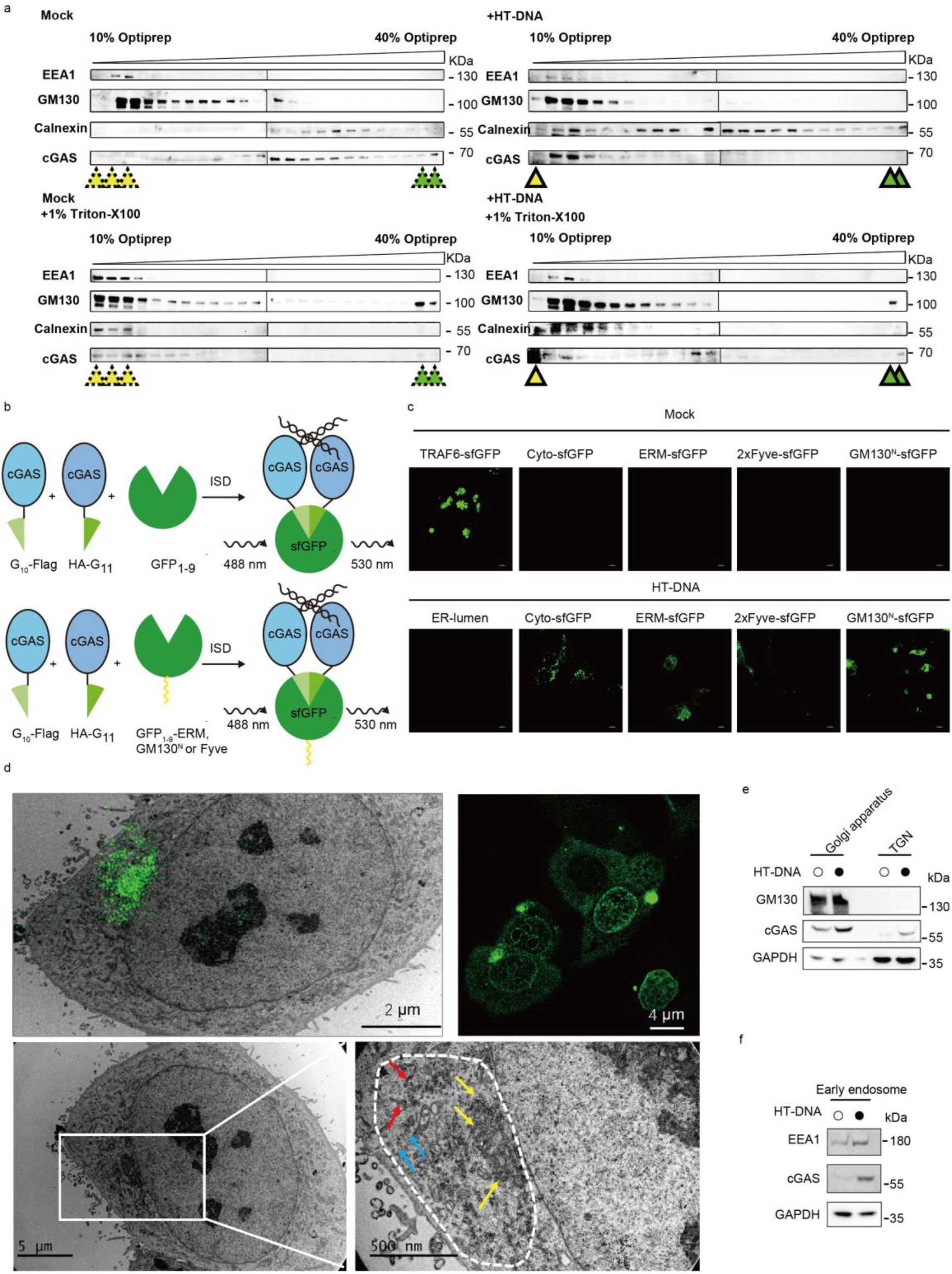
cGAS is associated with organelle membranes. (a) THP-1 cells were either untreated (Mock), stimulated with HT-DNA (2 μg/mL for 2 h), lysed in the presence of 1% Triton X-100, or subjected to both HT-DNA stimulation and Triton X-100 lysis, as indicated. Cytoplasmic extracts were subjected to OptiPrep density gradient centrifugation (10%–40%). Twenty-three fractions were collected from top (light) to bottom (dense) and analyzed by immunoblotting using antibodies against EEA1 (early endosome marker), GM130 (Golgi marker), Calnexin (endoplasmic reticulum marker), and cGAS. The yellow dashed triangle indicates that the distribution of the endoplasmic reticulum is affected by Triton X-100, while the yellow solid triangle indicates that cGAS remains present in the redistributed ER fractions. The green dashed triangle marks the redistribution of the Golgi apparatus upon Triton X-100 treatment, and the green solid triangle shows that cGAS is still retained in the altered Golgi fractions. (b) Schematic diagram showing the experimental setup for detecting cGAS interaction and activation using the split sfGFP system. cGAS is tagged with G_10_-Flag and HA-G_11_, which in the presence of DNA, bring together GFP_1−9_, G_10_ and G_11_ fragments to reconstitute sfGFP, allowing fluorescence detection. Variants include cytosolic sfGFP, ERM-sfGFP, 2xFyve-sfGFP and GM130^N^ for different subcellular localizations. (c) Fluorescence microscopy images showing the localization of sfGFP-tagged cGAS in different cellular compartments (cytosol, endoplasmic reticulum (ER), Golgi apparatus and endosome) with and without HT-DNA. For the positive control, sfGFP_1-9_ fragment was fused to TRAF6, a protein known to spontaneously form aggregates upon overexpression, thus enabling successful assembly of the full sfGFP. The negative control consisted of the human serum albumin signal peptide (MKWVTFISLLFLFSSAYSRGVFRR) fused to the N-terminus of sfGFP_1-9_ and a KDEL sequence fused to the C-terminus, ensuring retention in the ER lumen. Scale bars = 5 µm. (d) CLEM reveals the subcellular localization of cGAS to organelle membranes. Top-left: Correlative image combining light and electron microscopy, scale bar: 2 μm; Top-right: Confocal microscopy showing GFP-cGAS localization in HeLa cells stably expressing GFP-cGAS, following 2-hour transfection with HT-DNA (2 μg/ml), scale bar: 4 μm; Bottom-left: Transmission electron microscopy (TEM) image of the same cell, operating voltage: 100 kV, scale bar: 5 μm; Bottom-right: Enlarged view of the boxed region in the bottom-left panel, scale bar: 500 nm. The dashed white lines indicated the condensates of GFP-cGAS and HT-DNA. The red arrows indicate contacts between the endoplasmic reticulum (ER) and cGAS, the blue arrows indicate contacts between small vesicles and cGAS, and the yellow arrows indicate contacts between the Golgi apparatus and cGAS. (e-f) Fraction of Golgi apparatus (e) or early endosome (f) of THP1 cells in the absence or presence of HT-DNA (2 μg/ml). Subcellular fractionation of the Golgi apparatus (e) and early endosomes (f) was performed using commercial organelle isolation kits (Invent Biotechnologies) according to the manufacturer’s instructions. Western blot analysis was performed to detect the presence of cGAS in the purified fractions. GM130 and EEA1 were used as markers for the Golgi apparatus and early endosomes, respectively. GAPDH served as a cytosolic fraction control.

To further verify the subcellular localization of cGAS, we employed a split-GFP complementation assay. In this assay, cGAS was fused to one fragment of GFP (G_10_-Flag), and specific organelle markers were fused to another fragment of GFP (HA-G_11_) (Fig. 1b). The reconstitution of GFP fluorescence indicates the interaction and colocalization of cGAS with the respective organelle marker. We fused the self-organized TRAF6 to sfGFP as a positive control and the ER lumen localized sfGFP as a negative control. The strong GFP signal in the absence of HT-DNA and the disappeared GFP signal in the presence of HT-DNA indicates that the system is functional. We then found that the fluorescence was detected in cells expressing cytosolic sfGFP, ERM-sfGFP, GM130^N^-sfGFP and 2xFyve-sfGFP only in the presence of DNA, confirming that cGAS localizes to the cytosol, ER, Golgi apparatus and endosomes upon DNA stimulation (Fig. 1c).

To investigate the spatial relationship between activated cGAS and intracellular membranes, we performed correlative light and electron microscopy (CLEM) in HeLa cells stably expressing GFP-cGAS. Cells were transfected with HT-DNA to activate cGAS and induce cytoplasmic condensate formation. Confocal imaging revealed that GFP-cGAS formed distinct puncta in the cytosol and near the perinuclear region (Fig. 1d, top-right), suggestive of compartmentalization upon DNA stimulation. To further examine the membrane associations of these GFP-cGAS condensates, we used CLEM to overlay the fluorescence signal with ultrastructural context from TEM. As shown in the correlative image (Fig. 1d, top-left), GFP-cGAS aggregates localized to electron-dense cytoplasmic areas, adjacent to multiple membrane structures. Detailed electron microscopy of the same cell (Fig. 1d, bottom panels) revealed that the GFP-positive regions (outlined by dashed white lines) were in direct contact with several organelles. Specifically, we observed membrane contacts between cGAS and the endoplasmic reticulum (red arrows), small vesicular compartments resembling endosomes (blue arrows), and the stacked cisternae of the Golgi apparatus (yellow arrows). These observations indicate that, upon activation, cGAS forms biomolecular condensates that physically associate with multiple membrane-bound organelles, potentially influencing its activation and downstream signaling via membrane-dependent mechanisms.

To further substantiate our claims, we examine whether the subcellular localization of cGAS is altered in human monocytes THP1 cells. Golgi apparatus and early endosome fractions were isolated using commercial organelle-specific fractionation kits. Western blotting revealed that in steady state, cGAS was minimally detectable in the early endosomal and partially detectable in Golgi fractions. In contrast, when treated with HT-DNA, levels of cGAS showed markedly increased in both the GM130-positive Golgi fractions (Fig. 1e), and the EEA1-positive early endosomal fractions (Fig. 1f). Taken together, our results indicate that cGAS dynamically localizes to the ER, Golgi apparatus, and endosomes upon DNA stimulation.

### ZDHHC18 and MARCH8 facilitate organelle membrane localization of cGAS via IDR

In the previous paper, we reported that cGAS could be post-translationally modified by palmitoylation and ubiquitination, by ZDHHC18 on Golgi apparatus and MARCH8 on endosome, respectively (Shi et al., 2022b; Yang et al., 2022). We speculated that these two proteins were important for the localization of cGAS to Golgi apparatus and endosome. To facilitate the upcoming research on the commonalities of these two proteins, we name them membrane-bound proteins modulating cGAS activity (MEMCA). We found that MEMCA contain an intrinsically disordered region (IDR) at their N-terminus that points towards the cytoplasm (Fig. S1a and b). We speculated that MEMCA facilitate the localization of cGAS to organelle membranes via the IDR. To verify the hypothesis, we constructed HeLa cells stably co-expressing GFP-tagged cGAS and mCherry-tagged ZDHHC18 and stimulated them with Cy5-labeled interferon-stimulatory DNA (ISD). As shown in Fig. 2a, in the absence of Cy5-ISD, GFP-cGAS exhibited a diffuse cytoplasmic distribution, while mCherry-ZDHHC18 localized to distinct punctate structures. Upon Cy5-ISD stimulation, GFP-cGAS colocalized with mCherry-ZDHHC18, indicating that DNA stimulation promotes the interaction between cGAS and ZDHHC18 (Fig. 2b).

**Fig. 2.**
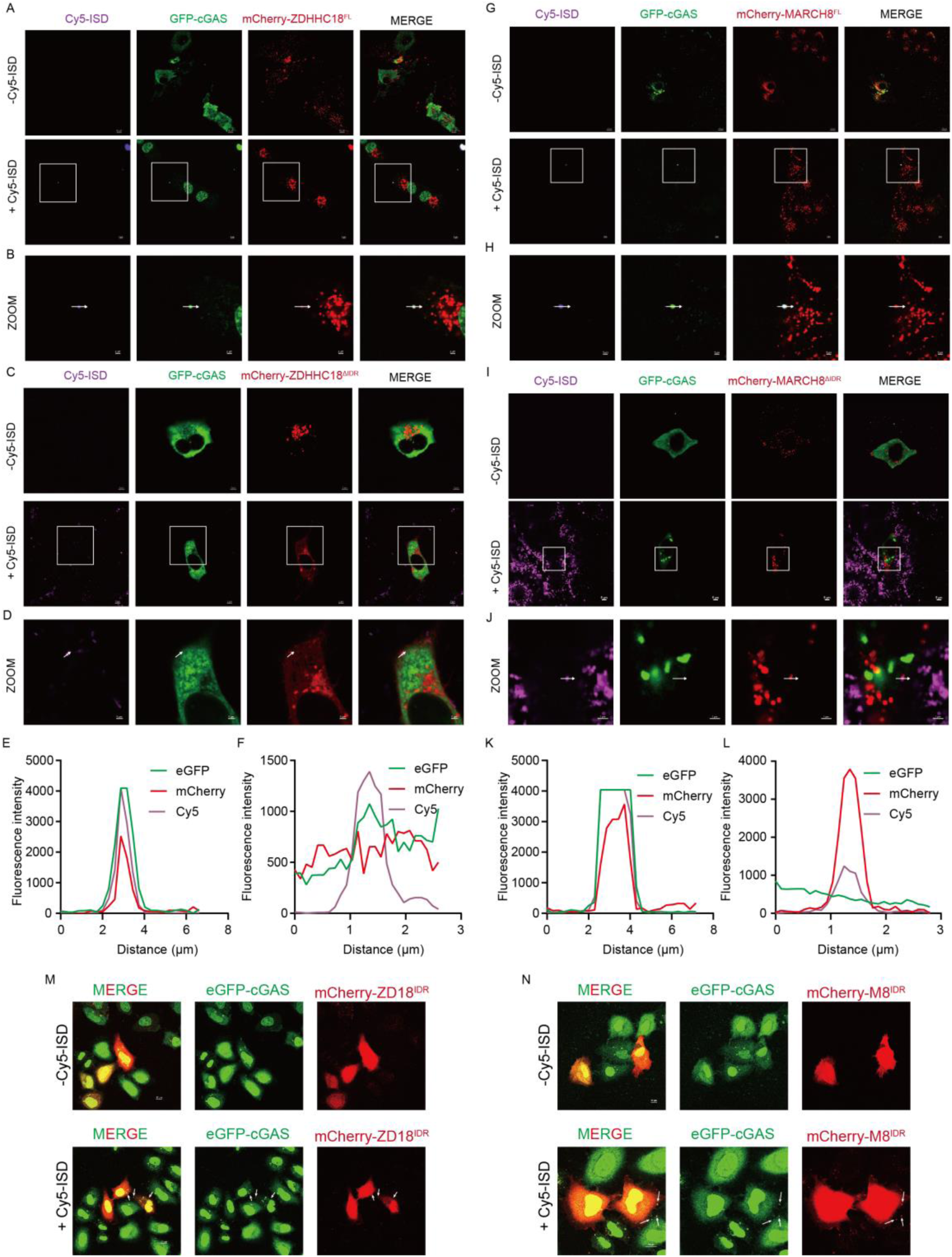
MEMCA facilitates organelle membrane localization of cGAS via IDR. (a) Representative confocal fluorescence microscopy images of HeLa cells stably expressing GFP-cGAS (green) and full-length mCherry-ZDHHC18^FL^ (red), transfected with or without Cy5-labeled interferon stimulatory DNA (Cy5-ISD, 2 μg/mL, 1 h; magenta). In the absence of DNA stimulation (–Cy5-ISD), GFP-cGAS is diffusely distributed in the cytosol and nucleus, while mCherry-ZDHHC18^FL^ displays punctate cytoplasmic localization with minimal overlap. Upon Cy5-ISD transfection (+Cy5-ISD), GFP-cGAS redistributes to cytoplasmic puncta that strongly co-localize with mCherry-ZDHHC18^FL^, suggesting condensate formation at DNA-associated compartments. Scale bars: 10 μm. (b) Enlarged high-magnification views of boxed regions from panel (a), highlighting the spatial convergence of GFP-cGAS, mCherry-ZDHHC18^FL^, and Cy5-ISD. White arrows indicate DNA-containing condensates that are positive for both cGAS and ZDHHC18^FL^, indicating a tripartite interaction upon activation. Scale bars: 2 μm. (c) Representative confocal images of HeLa cells stably expressing GFP-cGAS and the IDR deletion mutant mCherry-ZDHHC18^ΔIDR^. In unstimulated cells (–Cy5-ISD), both GFP-cGAS and ZDHHC18^ΔIDR^ show diffuse distributions, with no apparent co-localization. Upon Cy5-ISD transfection, GFP-cGAS forms DNA-associated puncta, but ZDHHC18^ΔIDR^ remains largely excluded from these structures, exhibiting weak or no co-localization. Scale bars: 10 μm. (d) High-magnification views of boxed regions from panel (c) reveal that although GFP-cGAS is recruited to Cy5-ISD-positive condensates (magenta and green), these structures lack enrichment of ZDHHC18^ΔIDR^ (red), suggesting that the IDR of ZDHHC18 is required for its DNA-induced co-localization with cGAS. Scale bars: 2 μm. (e) Line profile analysis of fluorescence intensity in (b). (f) Line profile analysis of fluorescence intensity in (d). (g) Representative confocal fluorescence microscopy images of HeLa cells stably expressing GFP-cGAS (green) and full-length mCherry-MARCH8^FL^ (red), transfected with or without Cy5-labeled interferon stimulatory DNA (Cy5-ISD, 2 μg/mL, 1 h; magenta). In the absence of DNA stimulation (–Cy5-ISD), GFP-cGAS is diffusely distributed in the cytosol and nucleus, while mCherry-MARCH8^FL^ displays punctate cytoplasmic localization with minimal overlap. Upon Cy5-ISD transfection (+Cy5-ISD), GFP-cGAS redistributes to cytoplasmic puncta that strongly co-localize with mCherry-MARCH8^FL^, suggesting condensate formation at DNA-associated compartments. Scale bars: 10 μm. (h) Enlarged high-magnification views of boxed regions from panel (g), highlighting the spatial convergence of GFP-cGAS, mCherry-MARCH8^FL^, and Cy5-ISD. White arrows indicate DNA-containing condensates that are positive for both cGAS and MARCH8^FL^, indicating a tripartite interaction upon activatio. Scale bars=2 µm. (i) Representative confocal images of HeLa cells stably expressing GFP-cGAS and the IDR deletion mutant mCherry-MARCH8^ΔIDR^. In unstimulated cells (–Cy5-ISD), both GFP-cGAS and MARCH8^ΔIDR^ show diffuse distributions, with no apparent co-localization. Upon Cy5-ISD transfection, GFP-cGAS forms DNA-associated puncta, but MARCH8^ΔIDR^ remains largely excluded from these structures, exhibiting weak or no co-localization. Scale bars=10 µm. (j) High-magnification views of boxed regions from panel (c) reveal that although GFP-cGAS is recruited to Cy5-ISD-positive condensates (magenta and green), these structures lack enrichment of MARCH8^ΔIDR^ (red), suggesting that the IDR of MARCH8 is required for its DNA-induced co-localization with cGAS. Scale bars=2 µm. (k) Line profile analysis of fluorescence intensity in (h). (l) Line profile analysis of fluorescence intensity in (j). (m and n) HeLa cells stably co-expressing eGFP-cGAS (green) and mCherry-ZDHHC18^IDR^ (red; IDR-only fragment of ZDHHC18) (m) and mCherry-MARCH8^IDR^ (red; IDR-only fragment of MARCH8) (n) were transfected with or without Cy5-labeled interferon stimulatory DNA (Cy5-ISD, 2 μg/mL, magenta) for 1 hour, followed by fixation and confocal microscopy. Each panel shows the merged image (left), eGFP-cGAS signal (middle), and mCherry signal (right). Scale bar: 10 μm.

Next, we investigated the role of the IDR of ZDHHC18 in this interaction. We generated a mutant of ZDHHC18 lacking its IDR (ZDHHC18^ΔIDR^) and construct HeLa cells stably co-expressing ZDHHC18^ΔIDR^ and GFP-cGAS. As shown in Fig. 2c, in the absence of Cy5-ISD, GFP-cGAS and mCherry-ZDHHC18^ΔIDR^ exhibited a diffuse distribution. Upon Cy5-ISD stimulation, GFP-cGAS did not colocalize with mCherry-ZDHHC18^ΔIDR^, indicating that the IDR of ZDHHC18 is crucial for the DNA-stimulated interaction with cGAS (Fig. 2d).

We then quantify the degree of colocalization between GFP-cGAS, mCherry-ZDHHC18, and Cy5-ISD. The fluorescence intensity profiles showed a significant overlap between GFP-cGAS and mCherry-ZDHHC18 in the presence of Cy5-ISD, while no significant overlap was observed with ZDHHC18^ΔIDR^ (Fig. 2e and f).

Similar results were obtained when we explored the interaction between cGAS and MARCH8. In the absence of DNA, both mCherry-MARCH8 and the mutant of MARCH8 lacking its IDR (MARCH8^ΔIDR^) showed no interaction with GFP cGAS. However, in the presence of Cy5-ISD, mCherry-MARCH8, but not MARCH8^ΔIDR^, exhibited a punctate distribution with cGAS in the presence of Cy5-ISD (Fig. 2g to j). Fluorescence intensity profiles also showed significant overlap between GFP-cGAS and mCherry-MARCH8 in the presence of Cy5-ISD, but not MARCH8^ΔIDR^ (Fig. 2k and l).

To further confirm the role of IDR of ZDHHC18 or MARCH8 in the association with cGAS, we examined the spatial distribution of cGAS in HeLa cells stably co-expressing either ZDHHC18^IDR^ or MARCH8^IDR^ mutants. Fig. 2m and n show that in the absence of Cy5-ISD, GFP-cGAS colocalize with either mCherry-ZDHHC18^IDR^ or mCherry-MARCH8^IDR^ in diffuse cytoplasm. In the presence of Cy5-ISD, GFP-cGAS, mCherry-ZDHHC18^IDR^ or mCherry-MARCH8^IDR^ colocalized in puncta induced by ISD, indicating the association between the IDR of ZDHHC18 or MARCH8, cGAS and dsDNA. In summary, our results demonstrate that ZDHHC18 and MARCH8 play a crucial role in the subcellular localization of cGAS to organelle membranes in response to DNA stimulation. The IDRs of these proteins are vital for their interaction with cGAS, highlighting the importance of intrinsically disordered regions in modulating subcellular localization of cGAS.

### MEMCA^IDR^ forms liquid-liquid phase separation in cells

To clarify the molecular mechanism under which IDR of MEMCA modulating membrane association of cGAS, we analyzed the amino acids composition of ZDHHC18 and MARCH8. Both IDRs of ZDHHC18 and MARCH8 are mainly composed of hydrophobic amino acids including proline, serine and glycine (Fig. S1c and d). Given that IDR is often indispensable to form liquid-liquid phase separation in cells (Hyman et al., 2014), we speculated that MEMCA IDR forms biomolecular condensates to facilitate cGAS membrane association. To verify the hypothesis, we firstly investigate the role of MEMCA IDR in the formation of biomolecular condensates.

First, we constructed optogenetic constructs of ZDHHC18 and MARCH8 by fusing their IDRs to mCherry and the blue light-sensitive Cry2 domain (Fig. 3a). These constructs allowed us to control the oligomerization and phase separation of ZDHHC18 and MARCH8 using blue light (488 nm). As shown in Fig. 3b and c, opto-MARCH8^IDR^ and opto-ZDHHC18^IDR^ showed diffuse distribution in the absence of light. Upon blue light exposure, mCherry-labeled opto-MARCH8^IDR^ and opto-ZDHHC18^IDR^ appears puncta, indicating biomolecular condensation forms.

**Fig. 3.**
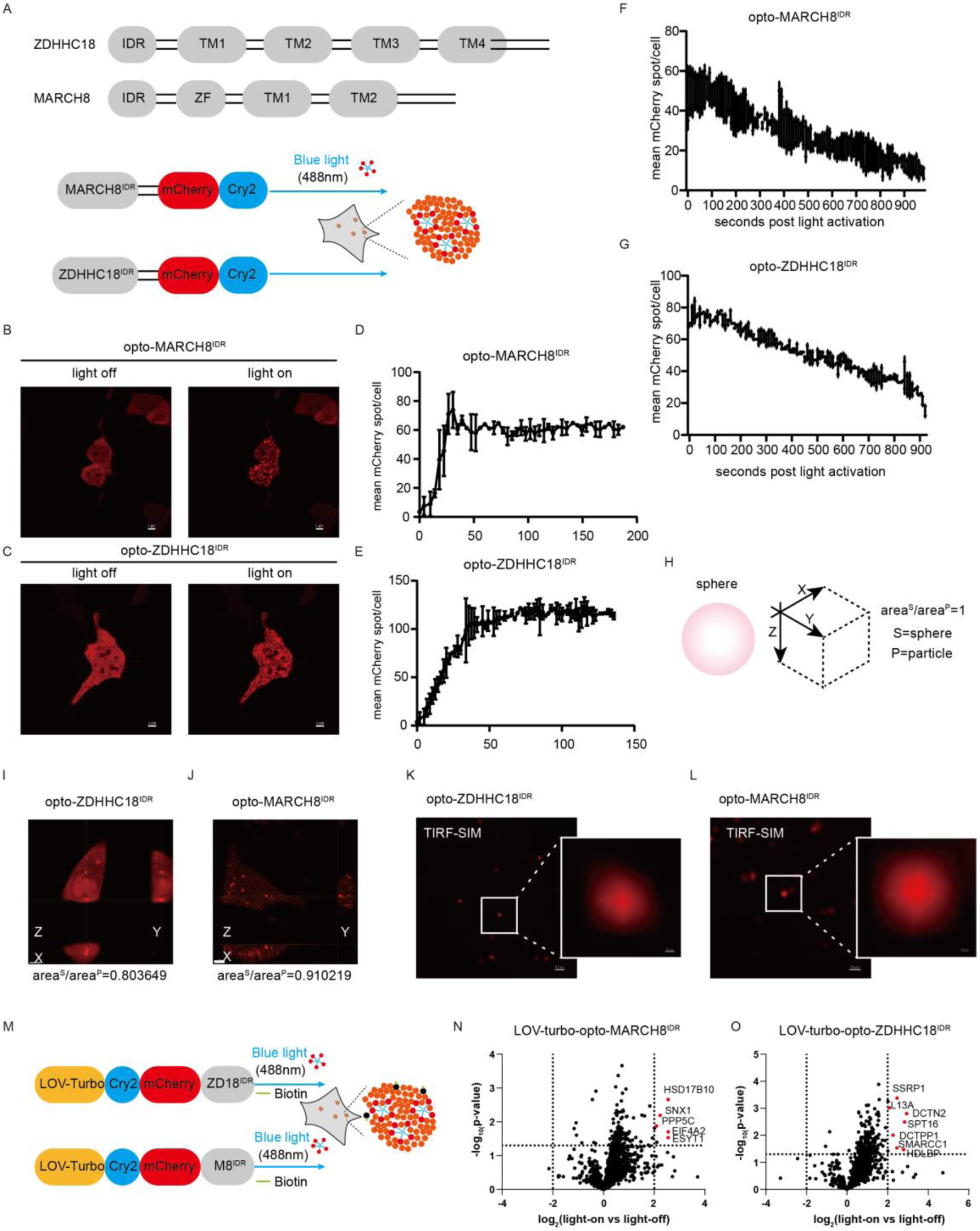
MEMCA^IDR^ forms biomolecular condensates in cells. (a) Schematic diagram showing the structure of ZDHHC18 and MARCH8 proteins, highlighting their IDR and transmembrane domains (TM). The opto-droplets system utilizes Cry2-mCherry fused to the IDR of MARCH8 and ZDHHC18 to induce condensate formation upon blue light (488 nm) activation. (b) Fluorescence microscopy images of cells expressing opto-MARCH8^IDR^ with and without blue light activation. The light-induced condensate formation is shown in the presence of blue light. Light ON: 3 min cycles of 4s light (488 nm) and 10s resting. Representative fluorescence images are shown. Scale bars=5 µm. (c) Fluorescence microscopy images of cells expressing opto-ZDHHC18^IDR^ with and without blue light activation. The light-induced condensate formation is shown in the presence of blue light. Light ON: 3 min cycles of 4s light (488 nm) and 10s resting. Representative fluorescence images are shown. Scale bars=5 µm. (d) Quantification of mean mCherry spot/cell for opto-MARCH8^IDR^ before and after light activation, showing the kinetics of condensate formation. (e) Quantification of mean mCherry spot/cell for opto-ZDHHC18^IDR^ before and after light activation, showing the kinetics of condensate formation. (f) After 5 minutes of light activation, blue light illumination was discontinued, and changes in the mCherry signal were subsequently monitored. Decay of mean mCherry spot/cell for opto-MARCH8^IDR^ after cessation of blue light activation, indicating the dynamics of condensate dissolution. (g) After 5 minutes of light activation, blue light illumination was discontinued, and changes in the mCherry signal were subsequently monitored. Decay of mean mCherry spot/cell for opto-ZDHHC18^IDR^ after cessation of blue light activation, indicating the dynamics of condensate dissolution. (h) Diagram illustrating the calculation of aspect ratio to determine the shape of the condensates. (i) Three-dimensional analysis of opto-ZDHHC18^IDR^ condensates showing the aspect ratio close to 1, indicating a spherical shape. Scale bar=5 µm. (j) Three-dimensional analysis of opto-MARCH8^IDR^ condensates showing the aspect ratio close to 1, indicating a spherical shape. Scale bar=5 µm. (k) TIRF-SIM images of opto-ZDHHC18^IDR^ condensates providing high-resolution details of the condensate structure. Scale bar=0.5 µm. (l) TIRF-SIM images of opto-MARCH8^IDR^ condensates providing high-resolution details of the condensate structure. Scale bar=0.5 µm. (m) Schematic diagram of the opto-IDR assay setup using LOV-Turbo and Cry2-mCherry to investigate the formation and composition of MEMCA^IDR^ condensates upon blue light activation. (n and o) Volcano plots showing proteins enriched in proximity to MARCH8^IDR^ (n) or ZDHHC18^IDR^ (o) following opto-genetically induced condensate formation using the LOV-Turbo system. THP-1 cells stably expressing LOV-Turbo-Cry2-mCherry-MARCH8^IDR^ or LOV-Turbo-Cry2-mCherry-ZDHHC18^IDR^ constructs. Cells were exposed to 488 nm blue light for 30 minutes (“light-on”) or maintained in the dark (“light-off”) in the presence of 500 μM biotin, followed by streptavidin pulldown and LC-MS/MS analysis of biotinylated proteins. Proteins are plotted by their log₂ fold-change (light-on vs light-off) on the x-axis and the –log₁₀(p-value) from unpaired Student’s t-tests on the y-axis. Vertical dotted lines represent ±1 log₂ fold change; horizontal dotted lines denote a significance threshold of p = 0.05. Significantly enriched proteins (right quadrant) are highlighted in red and labeled.

We quantified the dynamics of condensate formation by measuring the mean mCherry spot/cell over time. Fig. 3d and e show a rapid increase in the number of mCherry spots upon light activation, indicating the formation of condensates. This process reached a plateau within 150 seconds for both opto-MARCH8^IDR^ and opto-ZDHHC18^IDR^. However, the disassembly dynamics differed between the two proteins. Interestingly, upon cessation of blue light stimulation, the mean mCherry signal per cell gradually decreased in about 15 mins, indicating the disassembly of these condensates (Fig. 3f and g). Specifically, opto-MARCH8^IDR^ condensates disassembled more rapidly than opto-ZDHHC18^IDR^ condensates, which suggests distinct regulatory mechanisms for condensate dynamics. As a contrast, Cry2 oligomers disassemble spontaneously within 1–2 min (Shin et al., 2017). The enhanced stability of the droplets indicated the strong multivalent cooperative interactions within MEMCA IDR.

Changes in osmotic concentration influenced the formation of liquid-liquid phase separation in cells (Kilic et al., 2019). When we added sorbitol into cells to challenge the osmotic condition, the droplets stimulated by blue light disappeared (Fig. S1e and f). consistently, when treated with 1,6-HD, a widely used small molecular which disturbs hydrophobic interaction, blue light induced droplets also disappeared (Fig. S1e and f). These results indicated both electrostatic forces and hydrophobic interaction play important roles in the formation of MEMCA^IDR^ biomolecular condensates.

Liquid-liquid phase separation generally forms micrometer-sized biomolecular condensates. To dissect the biophysical feature of MEMCA^IDR^ condensates, we analyzed its aspect ratio using three-dimensional (3D) measurement. 3D reconstructions were performed in cells expressing opto-ZDHHC18^IDR^ and opto-MARCH8^IDR^, respectively. The area-to-sphere ratio was calculated to quantify the sphericity of the condensates. Opto-ZDHHC18^IDR^ condensates exhibited a lower area^s^/area^p^ ratio (0.803649), indicating less spherical and more irregular shapes compared to opto-MARCH8^IDR^ condensates (0.910219), which were more spherical. (Fig. 3h to j).

We then used Total Internal Reflection Fluorescence-Structured Illumination Microscopy (TIRF-SIM) to examine the nanoscale organization of opto-ZDHHC18^IDR^ and opto-MARCH8^IDR^ condensates (Fig. 3k and l). The high-resolution images revealed that these condensates possess a dense core surrounded by a less dense periphery, indicating a hierarchical organization of MEMCA condensations.

To identify potential interaction partners involved in the formation and regulation of these condensates, we performed proximity labeling assays using LOV-turbo-opto constructs of ZDHHC18^IDR^ and MARCH8^IDR^ (Fig. 3m). These constructs enable biotinylating of nearby proteins upon blue light activation. Mass spectrometry analysis revealed several candidates with significant enrichment in light-on versus light-off conditions (Fig. 3n and o). Notably, proteins involved in vesicular trafficking (e.g., SNX1, HSD17B10) and cytoskeletal regulation (e.g., PPP5C, ESRP1) were identified as interaction partners of MARCH8^IDR^. In contrast, proteins associated with chromatin remodeling (e.g., SSRP1, HDAC1) and transcriptional regulation (e.g., GPATCH1) were enriched for ZDHHC18^IDR^. In summary, our results demonstrate that the IDR of ZDHHC18 and MARCH8 can form dynamic biomolecular condensates in response to optogenetic activation. These condensates exhibit liquid-like properties and are enriched with specific interactors, implicating them in the regulation of membrane-associated signaling.

### MEMCA^IDR^ undergoes liquid-liquid phase separation in vitro

Purified IDR of proteins involved in the formation of biomolecular condensations could form liquid droplets in vitro (Lin et al., 2015; Ravindran et al., 2023; Sabari et al., 2018). To investigate the capacity of MEMCA^IDR^ to form phase-separated liquid droplets in vitro, we expressed and purified IDR of ZDHHC18 and MARCH8, respectively.

We first examined the phase separation behavior of GFP-tagged ZDHHC18^IDR^ and MARCH8^IDR^ at different protein concentrations and polyethylene glycol (PEG) concentrations. As shown in Fig. 4a, eGFP-ZDHHC18^IDR^ formed phase-separated droplets in the presence of increasing concentrations of PEG (5%, 10%, and 15%) and at various protein concentrations (5 μM, 10 μM, 25 μM, and 50 μM). The droplet formation was more pronounced at higher PEG and protein concentrations, indicating a concentration-dependent phase separation. Similarly, eGFP-MARCH8^IDR^ exhibited phase separation under the same conditions (Fig. 4b), with droplets forming at higher PEG and protein concentrations.

**Fig. 4.**
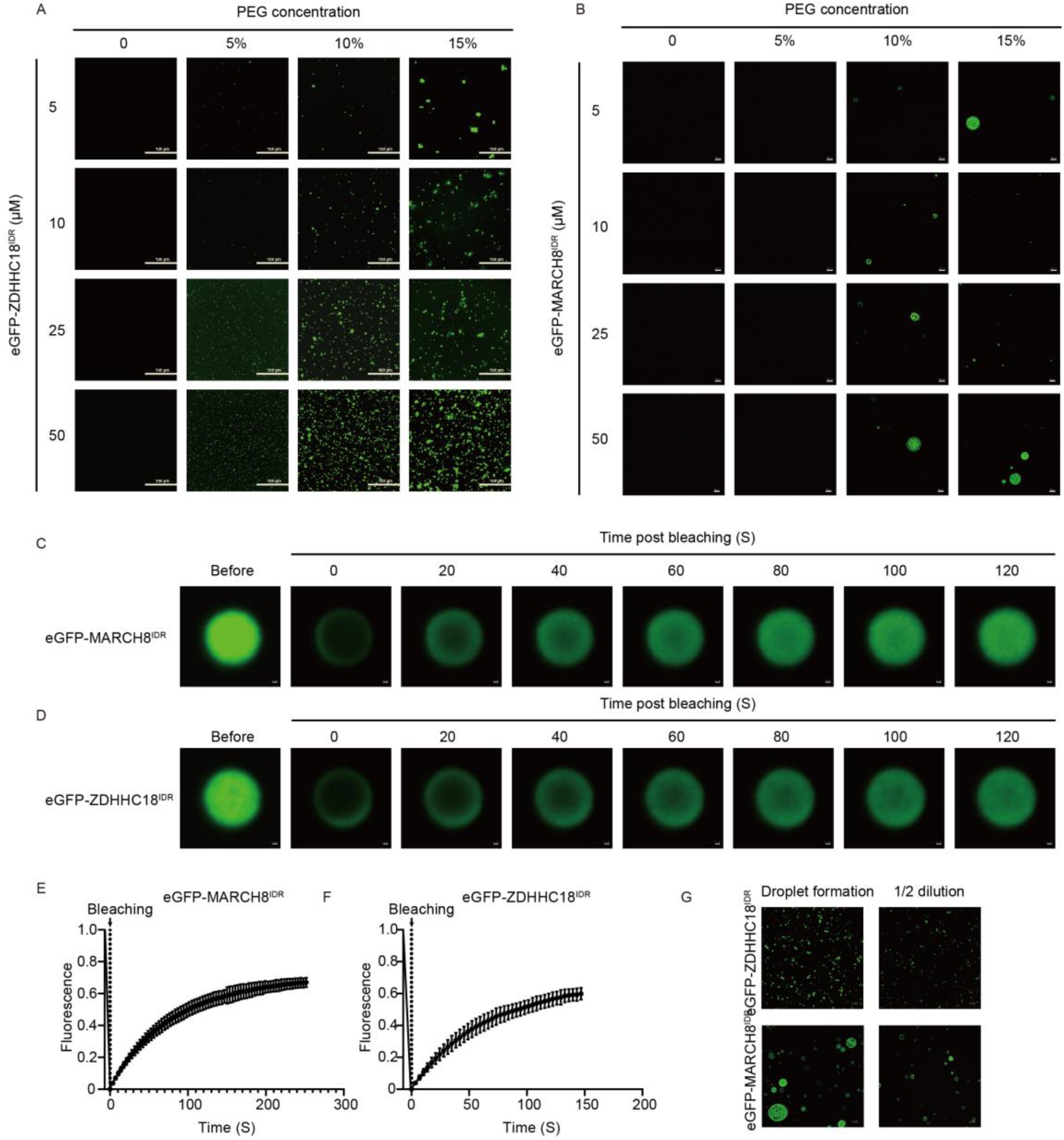
MEMCA^IDR^ undergoes liquid-liquid phase separation in vitro. (a) Phase separation of eGFP-ZDHHC18^IDR^ at varying concentrations (5, 10, 25, and 50 µM) and PEG-8000 concentrations (0, 5%, 10%, and 15%). Fluorescence microscopy images show the formation of liquid droplets indicating phase separation. Scale bars=100 µm. (b) Phase separation of eGFP-MARCH8^IDR^ at varying concentrations (5, 10, 25, and 50 µM) and PEG-8000 concentrations (0, 5%, 10%, and 15%). Fluorescence microscopy images show the formation of liquid droplets indicating phase separation. Scale bars=10 µm. (c) Fluorescence recovery after photobleaching (FRAP) analysis of eGFP-MARCH8^IDR^ (10µM) droplets with 10% PEG-8K. Images show the fluorescence recovery over time after photobleaching, indicating dynamic exchange within the droplets. Scale bar=0.2 µm. (d) Fluorescence recovery after photobleaching (FRAP) analysis of eGFP-ZDHHC18^IDR^ (10µM) droplets 10% PEG-8K. Images show the fluorescence recovery over time after photobleaching, indicating dynamic exchange within the droplets. Scale bar=0.2 µm. (e) Quantification of FRAP recovery curves for eGFP-MARCH8^IDR^ droplets, showing the mean fluorescence intensity over time post-bleaching. Time 0 indicates the end of photobleaching and the start of recovery. The mean ± SD are shown. N = 3 liquid droplets. (f) Quantification of FRAP recovery curves for eGFP-ZDHHC18^IDR^ droplets, showing the mean fluorescence intensity over time post-bleaching. Data represent the mean ± SEM from three independent experiments. Time 0 indicates the end of photobleaching and the start of recovery. The mean ± SD are shown. N = 3 liquid droplets. (g) Reversibility of droplet formation for eGFP-ZDHHC18^IDR^ (top row) and eGFP-MARCH8^IDR^ (bottom row) upon dilution. Fluorescence microscopy images show droplet formation at 25 µM protein concentration and the reduction in size and number of droplets upon 1/2 dilution. Scale bars = 20 µm.

We then investigated another crowding agents ficoll. Consistent with the result of PEG-8000, both ZDHHC18^IDR^ and MARCH8^IDR^ formed liquid droplets in ficoll (Fig. S2a and b), conforming phase separation of MEMCA^IDR^ in vitro.

To further characterize the dynamics of these phase-separated droplets, we performed fluorescence recovery after photobleaching (FRAP) experiments. Fig. 4c and d show representative images of FRAP recovery for eGFP-MARCH8^IDR^ and eGFP-ZDHHC18^IDR^ droplets, respectively. The fluorescence recovery over time indicates the dynamic nature of these droplets, with rapid recovery suggesting liquid-like properties.

Quantitative analysis of the FRAP recovery curves (Fig. 4e and f) revealed that eGFP-MARCH8^IDR^ droplets exhibited a faster recovery half-time compared to eGFP-ZDHHC18^IDR^ droplets. This suggests that MARCH8^IDR^ droplets are more dynamic and less stable than ZDHHC18^IDR^ droplets, which may reflect differences in the molecular interactions driving phase separation for these two proteins.

We next sought to test whether the droplets were irreversible aggregates or reversible phase-separated condensates. To do this, ZDHHC18^IDR^ and MARCH8^IDR^ were allowed to form droplets in an initial solution. Then the protein concentration was diluted by buffer with equal salt concentration. Upon dilution, the phase-separated droplets of both eGFP-ZDHHC18^IDR^ and eGFP-MARCH8^IDR^ dispersed, indicating that the phase separation process is reversible and dependent on the concentration of protein and crowding agents like PEG (Fig. 4g).

Weak interactions including electrostatic interactions and hydrophobic interactions are believed to drive the formation of liquid droplets. To rule out the biophysical properties of MEMCA^IDR^ droplets, we altered the salt concentrations of the buffer to determine the contribution of electrostatic interactions or added 1,6-hexanediol to determine the contribution of hydrophobic interactions in driving formation of the droplets. As shown in Fig. S2c, the size and numbers of both ZDHHC18^IDR^ and MARCH8^IDR^ forms decreased with increasing NaCl concentration (from 0 to 300 mM), indicating electrostatic interactions contribute to MEMCA^IDR^ droplets formation. We also found that treatment with 10% 1,6-hexanediol decreased the size of MEMCA^IDR^ droplets, indicating hydrophobic interactions also played important role in driving MEMCA^IDR^ droplets formation (Fig. S2d and e).

In conclusion, our study demonstrates that the IDRs of ZDHHC18 and MARCH8 undergo LLPS in vitro, forming dynamic and reversible phase-separated droplets.

### Phosphorylation regulates ZDHHC18^IDR^ condensation

Post-translational modification including phosphorylation regulates protein phase separation through altering electrostatic forces. We asked whether MEMCA condensation was regulated by phosphorylation. Specifically, we investigated how phosphorylation and phosphomimic mutations influence the formation and dynamics of biomolecular condensates.

To answer this question, we firstly detect phosphorylation on ZDHHC18 through LC-MS/MS analysis. We found ZDHHC18^IDR^ could be phosphorylated at S19, which was also reported by previous screening (Zhou et al., 2013), S30 and S366. We focused on phosphorylation of S19 and S30 which were in the ZDHHC18^IDR^ (Fig. S3 and Fig. S4a). We then investigated how phosphorylation and phosphomimic mutations influence the formation and dynamics of ZDHHC18^IDR^ biomolecular condensates.

To detect the influence of phosphorylation on MEMCA phase separation, we replaced these two serine (S) residues with aspartate (S→D) to mimic its phosphorylation. We also replaced serine with alanine (S→A) to mimic phosphorylation deficiency (Fig. S4a). These constructs were tagged with mCherry to visualize their localization and condensate formation. To investigate the effects of phosphorylation on phase separation, we expressed these constructs in cells and induced opto-droplets system.

As shown in Fig. S4b and c, the WT and S to D constructs rapidly formed punctate condensates upon light activation, while the S to A construct showed a delayed and less efficient condensate formation. This suggests that phosphorylation enhances the ability of ZDHHC18^IDR^ to undergo phase separation.

To further confirm this result, we purified mCherry-fused recombinant ZDHHC18^IDR^ protein with those mutations. We found that the WT and S to D constructs formed numerous condensates in the presence of 10% PEG, whereas the S to A construct exhibited fewer and smaller condensates. These results indicate that phosphorylation promotes the formation of larger and more numerous biomolecular condensates of ZDHHC18 (Fig. S4d).

To further assess the function of phosphorylation in formation process of ZDHHC18^IDR^ condensation, we firstly mixed GFP-ZDHHC18^IDR^ with PEG-8K to form phase separation (Fig. S4e and f). Addition of mCherry-ZDHHC18^IDR^ to pre-formed GFP-ZDHHC18^IDR^ condensates yielded yellow compartments, indicating that soluble molecules are recruited to ZDHHC18^IDR^ condensates. Surprisingly, we found that Addition of mCherry-ZDHHC18^IDR^ with S to A mutation significantly destroyed pre-formed GFP-ZDHHC18^IDR^ condensates, indicating a dominant negative effect of the phosphorylation deficient mutant in impairing ZDHHC18^IDR^ condensation formation. Taken together, these results suggested that phosphorylation regulates ZDHHC18 condensation (Fig. S4e and f). Taken together, these findings demonstrated that phosphorylation is important for ZDHHC18 condensation.

### MEMCA biomolecular condensation recruits cGAS and dsDNA

Given that MEMCA^IDR^ forms biomolecular condensates in cells and in vitro, we next explore its role in recruiting cGAS and dsDNA. We first examined the interaction between eGFP-cGAS and mCherry-tagged ZDHHC18^IDR^ or MARCH8^IDR^ under various conditions, including buffer alone, the presence of 1,6-hexanediol (1,6-HD), and high salt concentration. As shown in Fig. 5a and b, eGFP-cGAS and mCherry-ZDHHC18^IDR^ or mCherry-MARCH8^IDR^ formed distinct condensates in buffer conditions. However, the presence of 1,6-HD, a known disruptor of weak hydrophobic interactions, significantly reduced droplet formation, indicating the importance of hydrophobic interactions in maintaining these condensates. High salt conditions, which disrupt electrostatic interactions, also diminished droplet formation, suggesting that both hydrophobic and electrostatic interactions are crucial for the stability of these biomolecular condensates (Fig. 5a and b).

**Fig. 5.**
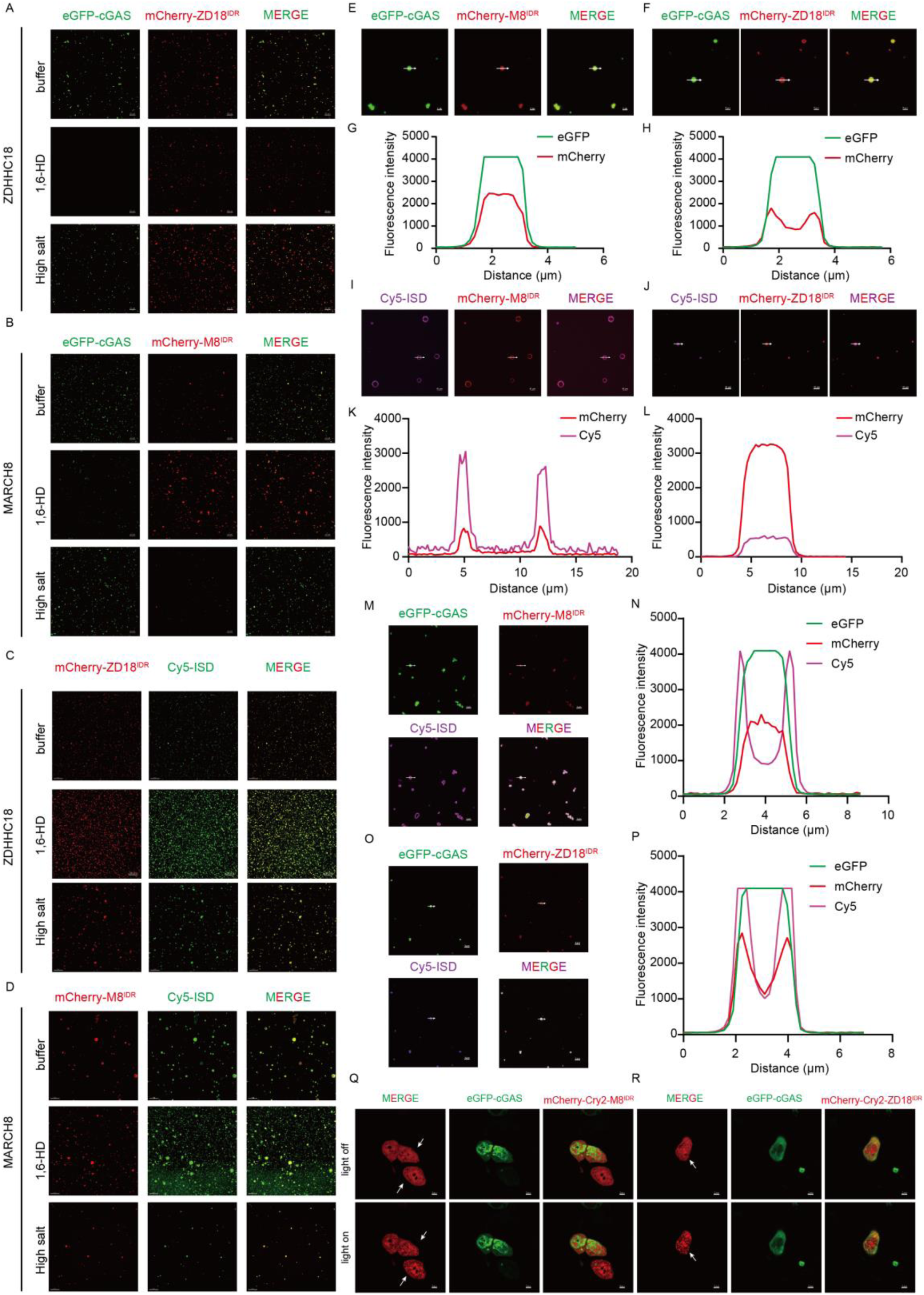
MEMCA^IDR^ biomolecular condensates recruit cGAS and dsDNA. (a) Fluorescence microscopy images showing the recruitment of eGFP-cGAS (green) (2 μM) to mCherry-ZDHHC18^IDR^ (red) (10 μM) condensates with treatment of 16-HD or high salt (300 mM NaCl). The images demonstrate the effect of these treatments on the recruitment process. Scale bars=20 µm. (b) Fluorescence microscopy images showing the recruitment of eGFP-cGAS (green) (2 μM) to mCherry-MARCH8^IDR^ (red) (10 μM) condensates with treatment of 16-HD or high salt (300 mM NaCl). The images demonstrate the effect of these treatments on the recruitment process. Scale bars=20 µm. (c) Fluorescence microscopy images showing the recruitment of Cy5-ISD (green) (0.5 μM) to mCherry-ZDHHC18^IDR^ (red) (10 μM) condensates with treatment of 16-HD or high salt (300 mM NaCl). The images demonstrate the effect of these treatments on the recruitment process. Scale bars=20 µm. (d) Fluorescence microscopy images showing the recruitment of Cy5-ISD (green) (0.5 μM) to mCherry-MARCH8^IDR^ (red) (10 μM) condensates under different conditions: buffer, 16-HD, and high salt. The images demonstrate the effect of these treatments on the recruitment process. Scale bars=20 µm. (e and f) Fluorescence microscopy images showing in vitro condensates formed by recombinant eGFP-cGAS (green, 2 μM) and mCherry-MARCH8^IDR^ (red, 10 μM) (e) or mCherry-ZDHHC18^IDR^ (red, 10 μM) (f) incubated in reaction buffer (20 mM Tris pH 7.5, 150 mM NaCl, 5 mM MgCl_2_) at room temperature for 30 minutes. Confocal images show distinct phase-separated droplets containing both fluorescent proteins. Arrows indicate representative condensates used for fluorescence intensity line profiling. Scale bars: 5 μm. (g and h) Fluorescence intensity profiles along the white line in panel e (g) and f (h), plotted as a function of distance (μm). Fluorescence intensities were extracted using ImageJ. (i and j) Fluorescence microscopy images showing the result of in vitro mixing of Cy5-labeled interferon stimulatory DNA (Cy5-ISD, magenta, 0.5 μM) with recombinant mCherry-MARCH8^IDR^ (red, 10 μM) (i) or mCherry-ZDHHC18^IDR^ (red, 10 μM) (j) in reaction buffer (20 mM Tris pH 7.5, 150 mM NaCl, 5 mM MgCl_2_). Samples were incubated at room temperature for 30 minutes prior to imaging by confocal microscopy. White arrows indicate representative puncta. Scale bars: 5 μm. (k and l) Fluorescence intensity line-scan analysis along the white arrow shown in i (k) and j (l), plotting Cy5 (magenta) and mCherry (red) signal intensities across the selected condensates. Fluorescence intensities were extracted using ImageJ. (m and o) Representative fluorescence microscopy images showing in vitro reconstitution of phase-separated condensates formed by mixing recombinant eGFP-cGAS (green, 2 μM), mCherry-MARCH8^IDR^ (red, 10 μM) (m) or mCherry-ZDHHC18^IDR^ (red, 10 μM) (o), and Cy5-labeled interferon stimulatory DNA (Cy5-ISD, magenta, 0.5 μM) in reaction buffer (20 mM Tris pH 7.5, 150 mM NaCl, 5 mM MgCl_2_). Following 30 minutes of incubation at room temperature, samples were imaged using confocal microscopy. The merged image shows clear overlap of all three components within the same condensates. White arrows indicate regions of interest used for subsequent line profile analysis. Scale bars: 5 μm. (n and p) Quantification of fluorescence intensity along the white line indicated in (m) and (o). Line-scan analysis reveals co-enrichment of eGFP-cGAS (green), mCherry-MARCH8^IDR^ (m) or ZDHHC18^IDR^ (o) (red), and Cy5-ISD (magenta) within the same micron-scale condensates. Fluorescence intensities were extracted using ImageJ. (q and r) HeLa cells stably expressing eGFP-cGAS (green) and mCherry-Cry2-MARCH8^IDR^ (red) (q) or mCherry-Cry2-ZDHHC18^IDR^ (red) (r) were subjected to light-inducible phase separation. Cells were maintained in the dark (“light off”, top row) or exposed to 488 nm blue light for 2 minutes (“light on”, bottom row), followed by confocal imaging. The white arrows indicate the cells subjected to light activation. Each panel shows merged (left), green (middle), and red (right) fluorescence channels. Scale bars=5 µm.

We then replaced cGAS with its ligand, Cy5-labelled ISD, in the recruitment experiment. ISD was also present in both mCherry-ZDHHC18^IDR^ or mCherry-MARCH8^IDR^ droplets, indicating DNA could also be recruited into MEMCA condensates (Fig. 5c and d). Specifically, 1,6-HD treatment significantly promoted DNA recruitment of mCherry-ZDHHC18^IDR^ or mCherry-MARCH8^IDR^. Since DNA could not form liquid droplets alone in the presence of PEG-8K or 1,6-HD, we concluded that MEMCA^IDR^ recruited DNA through weak interaction.

To further confirm the recruitment of cGAS into these condensates, we performed colocalization studies. Fig. 5e and f show that eGFP-cGAS colocalized with mCherry-MARCH8^IDR^ and mCherry-ZDHHC18^IDR^ condensates, respectively. The fluorescence intensity profiles in Fig. 5g and h demonstrate a significant overlap between eGFP and mCherry signals, confirming the sequestration of cGAS into the condensates formed by the IDR of ZDHHC18 and MARCH8.

Next, we investigated the ability of these condensates to recruit dsDNA by using Cy5-labeled ISD (Cy5-ISD). As shown in Fig. 5i to l, mCherry-MARCH8^IDR^ and mCherry-ZDHHC18^IDR^ condensates effectively recruited Cy5-ISD, as evidenced by the colocalization of the Cy5 signal with the mCherry-labeled droplets. indicating that these condensates can sequester both cGAS and dsDNA. As a contrast, no liquid droplets were observed in Cy5-ISD with PEG or 1.6-HD, conforming the recruitment of Cy5-ISD into MEMCA^IDR^ droplets (Fig. S5a). Moreover, we concluded that the interaction between MEMCA and DNA were relatively weak, because there is no direct interaction between MARCH8^IDR^ or ZDHHC18^IDR^ with Cy5-ISD (Fig. S5b).

We then examined the combined recruitment of cGAS and dsDNA into these condensates. As shown in Fig. 5m to p, eGFP-cGAS colocalized with mCherry-MARCH8^IDR^ or mCherry-ZDHHC18^IDR^, and Cy5-ISD, demonstrating that the IDR of ZDHHC18 and MARCH8 can form condensates that simultaneously recruit cGAS and dsDNA.

Next, we explored which one is the driving force between cGAS and MEMCA. To do so, we used light-activated optogenetic constructs of ZDHHC18^IDR^ and MARCH8^IDR^ fused to Cry2. In the absence of blue light stimulation, both cGAS and MEMCA^IDR^ were diffuse, indicating a steady state. Upon blue light exposure, mCherry-Cry2-ZDHHC18^IDR^ or mCherry-Cry2-MARCH8^IDR^ forms phase-separated droplets in cells, whereas eGFP-cGAS keeps diffuse state, indicating biomolecular condensation of MEMCA^IDR^ could not recruit cGAS alone (Fig. 5q and r). On the contrary, when we examine the concentration diagram of MEMCA with cGAS in LLPS in vitro, we found that the concentration of cGAS was rather important than that of both ZDHHC18^IDR^ or MARCH8^IDR^, suggesting that cGAS mainly drives the recruitment of MEMCA (Fig. S6a and b).

In summary, our results demonstrate that the IDR of ZDHHC18 and MARCH8 facilitates the formation of biomolecular condensates that can sequester cGAS and dsDNA.

### Interaction between cGAS and MEMCA^IDR^ is dependent on DNA

To further investigate the interaction between MEMCA^IDR,^ cGAS, and DNA, we performed GST-pulldown experiments. The results showed that MARCH8^IDR^/ZDHHC18^IDR^ could capture cGAS only in the presence of DNA (Fig. S7a to d). Sequence analysis revealed that MARCH8^IDR^/ZDHHC18^IDR^ contains numerous positively and negatively charged amino acids, which we hypothesized to be critical for their binding to the cGAS-DNA complex (Fig. S1c and d). To test this, we mutated all these charged amino acids to alanine and performed GST-pulldown experiments. The mutants were unable to bind the cGAS-DNA complex, suggesting that MARCH8^IDR^/ZDHHC18^IDR^ recruits the cGAS-DNA complex through these charged amino acids (Fig. S7a to d). Consistent results were obtained using mouse cGAS in GST-pulldown experiments, indicating that this mechanism is conserved between humans and mice (Fig. S7a to d).

To assess the affinity between MEMCA^IDR^, cGAS, and DNA, we conducted SPR experiments. The results indicated that MARCH8^IDR^/ZDHHC18^IDR^ did not bind to DNA or cGAS individually but exhibited micromolar-level affinity (K_D_ approximately 0.4 µM) for the cGAS-DNA complex (Fig. S7e and f).

Taken together, these results indicated that MEMCA^IDR^ interacts with cGAS in a DNA-dependent manner.

### MEMCA condensation regulates cGAS activity

To investigate the biological role of MEMCA^IDR^-mediated recruitment of cGAS to organelle membranes, we utilized a rapamycin-inducible dimerization system to induce the interaction between cGAS-FRB and FKBP12 or the organelle membrane associated ERM-mCherry-FKBP12, mCherry-FKBP12-2xFYVE or GM130^N^-mCherry-FKBP12. In the presence of rapamycin, ERM-mCherry-FKBP12, mCherry-FKBP12-2xFYVE or GM130^N^-mCherry-FKBP12 would recruit cGAS to localized to ER, endosome or Golgi apparatus, respectively. Whereas the FKBP12 group did not alter distribution of cytosolic cGAS (Fig. 6a). To detect the interference of membrane recruitment by MEMCA^IDR^ in cGAS enzymatic activity, we constructed stable THP-1 cell line with expression of cGAS-FRB and the FKBP12 constructs and measured cGAMP production by cGAMP bioassay with or without rapamycin (Fig. S8). The results showed that in the presence of HT-DNA, cGAMP production was not influenced in the absence of rapamycin, whereas the ERM-mCherry-FKBP12 or mCherry-FKBP12-2xFYVE group showed a significant reduction in cGAMP production in the presence of rapamycin, indicating that targeting cGAS to ER or endosome reduced cGAS activity (Fig. S8). Similar results were observed in the GM130^N^-mCherry-FKBP12 group (Fig. 6b). The results indicate that the membrane association significantly reduced enzymatic activity of cGAS. To further confirm whether this inhibitory effect is caused by ZDHHC18 or MARCH8, we knocked out ZDHHC18 or MARCH8 in THP-1 cells. The results showed that the knockout of ZDHHC18 or MARCH8 relieved the inhibitory effect of the Golgi and endosomes on cGAS (Fig. 6b and c). This indicates that ZDHHC18 and MARCH8 are important for the regulation of cGAS localized on these organelles.

**Fig. 6.**
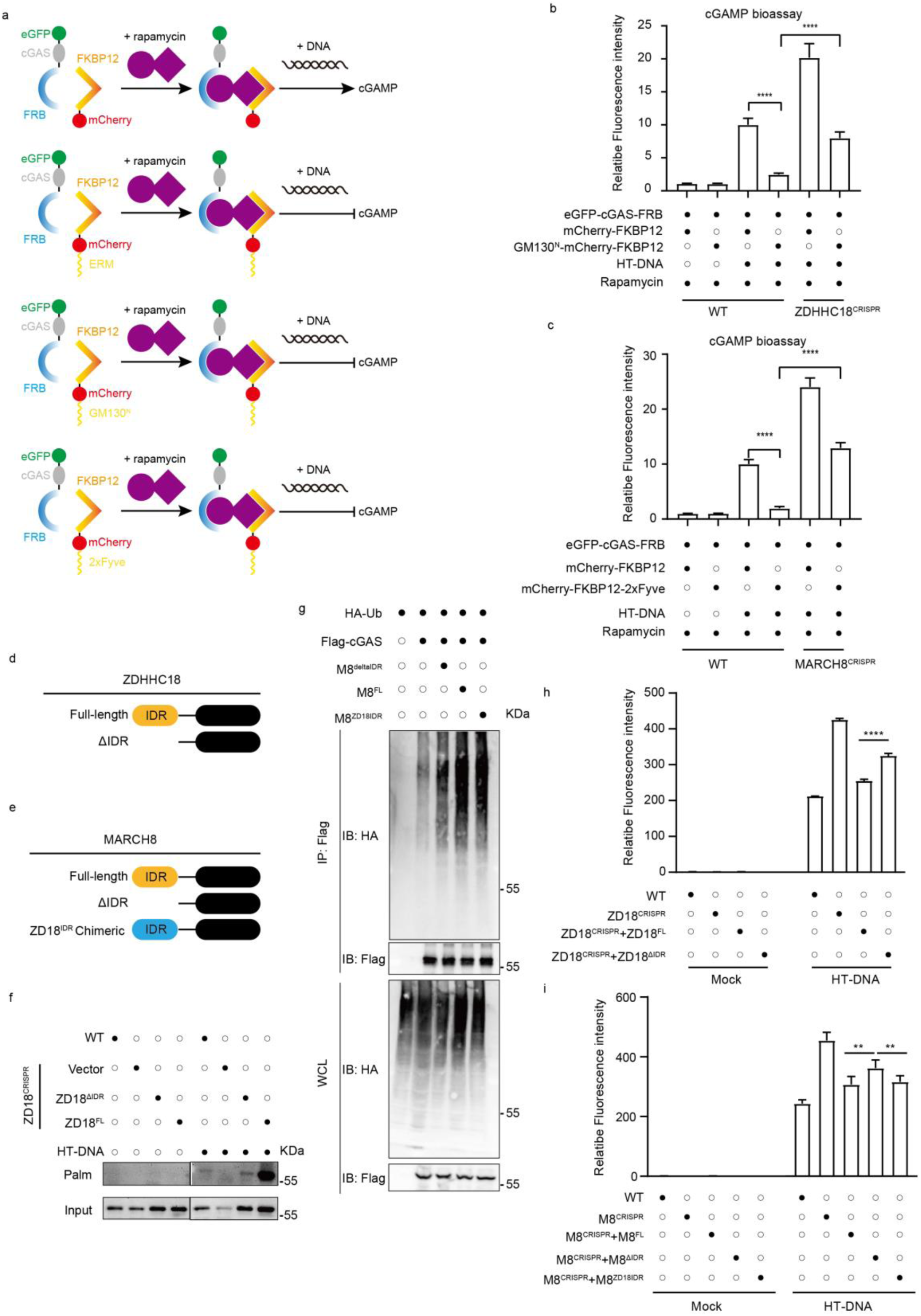
MEMCA biomolecular condensates regulates cGAS activity. (a) Schematic representation of the experimental setup to study cGAS activity regulation via rapamycin-induced biomolecular condensates. Fusion proteins eGFP-cGAS-FRB and mCherry-FKBP12 or its variants (ERM-mCherry-FKBP12, mCherry-FKBP12-2xFYVE and GM130^N^-mCherry-FKBP12) were stably expressed in THP-1 cells via lenti-virus package. Addition of rapamycin induces dimerization, leading to the formation of biomolecular condensates, followed by HT-DNA (2 μg/mL) stimulation for 6 hours to measure cGAS activity. (b-c) THP-1 cells or CRISPR-mediated knockout lines for ZDHHC18 (b) and MARCH8 (c) were stably transduced with eGFP-cGAS-FRB and various mCherry-FKBP12 fusion constructs (GM130^N^-mCherry-FKBP12 (b) and mCherry-FKBP12-2xFYVE (c)) to direct cGAS to specific organelles upon rapamycin treatment. Cells were treated with 100 nM rapamycin for 30 minutes to induce FRB-FKBP12 dimerization, followed by stimulation with 2 μg/mL HT-DNA for 6 hours. Intracellular cGAMP levels were quantified using a luciferase-based bioassay in THP-1 Lucia™ ISG cells (InvivoGen), and data are presented as relative fluorescence intensity. Data are presented as mean ± SEM; **** P<0.0001. (d) Schematic of ZDHHC18 constructs used in (f): full-length ZDHHC18 and ΔIDR ZDHHC18 lacking the intrinsically disordered region. (e) Schematic of MARCH8 constructs used in (g): full-length MARCH8 and ZD18^IDR^ chimeric construct where MARCH8^IDR^ is replaced with ZDHHC18^IDR^. (f) Western blot analysis of ZDHHC18^CRISPR^ and ZDHHC18 reconstituted cells showing the levels of palmitoylated (Palm) and input proteins in the presence and absence of HT-DNA. (g) Cells were co-transfected with HA-ubiquitin, Flag-cGAS, and either MARCH8 variants. Immunoprecipitation (IP) using anti-Flag antibodies followed by immunoblotting (IB) with anti-HA antibodies was performed to detect ubiquitinated cGAS. Whole cell lysates (WCL) were also probed with anti-Flag and anti-HA antibodies to confirm expression levels. (h and i) THP-1 Lucia™ ISG reporter cells were used to assess interferon pathway activity upon cytosolic DNA stimulation. CRISPR-Cas9-mediated knockout cell lines were generated for ZDHHC18 (h) or MARCH8 (i), and complemented with full-length, intrinsically disordered region–deleted (ΔIDR), or IDR-truncated constructs, as indicated. Cells were transfected with HT-DNA (2 μg/mL) or mock treated for 18 hours, and luciferase activity was measured as a readout of ISG pathway activation. Bar graphs show relative fluorescence intensity (mean ± SEM) from four biological replicates. Data are presented as mean ± SEM; **** P<0.0001. ** P<0.01.

Next, we examined the specific contributions of the IDR of ZDHHC18 and MARCH8 to this regulatory mechanism. We generated constructs with full-length ZDHHC18 and MARCH8, as well as constructs lacking the IDR (ΔIDR) and a specific chimeric MARCH8 constructs where the MARCH8^IDR^ were substituted by ZDHHC18^IDR^ (Fig. 6d and e). We asked whether the alteration in IDR influence the ability of MEMCA to catalyze cGAS PTM by employing these constructs in palmitoylation assay or ubiquitination assay. we found that cGAS palmitoylation was depleted in ZDHHC18 knockout THP1 cells in the presence of DNA, which was consistent with the previous report (Shi et al., 2022b). However, reconstituted full-length ZDHHC18, but not the IDR deleted ZDHHC18, restored cGAS palmitoylation level, indicating IDR was important for ZDHHC18 to catalyze cGAS palmitoylation (Fig. 6f). We then detected cGAS ubiquitination catalyzed by MARCH8. As shown in Fig. 6g, the full-length MARCH8 and chimeric construct containing ZDHHC18^IDR^ were more effective in promoting cGAS ubiquitination, compared to the ΔIDR constructs, indicating that the presence of IDR is critical for facilitating cGAS ubiquitination catalyzed by MARCH8 (Fig. 6g).

We further quantified the effect of IDR in modulating cGAMP production by reconstituting those constructs in THP1 lucia ISG reporter cells. The result showed that both MARCH8 and ZDHHC18 knockout enhanced luciferase expression in the presence of DNA, whereas reconstitution of full-length, but not IDR-deleted constructs, rescued the inhibition effect of both MARCH8 and ZDHCH18 (Fig. 6h and i). Notably, the chimeric constructs of MARCH8 with swapped ZDHHC18^IDR^ retained the ability to inhibit cGAS activity, highlighting the importance of IDR in the regulatory process (Fig. 6h and i). Taken together, these results suggested that MEMCA condensation modulated cGAS activity by recruiting cGAS to organelle membranes.

In summary, we believe we have uncovered the following process: In its resting state without DNA stimulation, cytoplasmic cGAS does not interact with membrane proteins on organelles; Upon sensing cytoplasmic double-stranded DNA, cGAS binds to DNA and undergoes phase separation, initiating downstream innate immune signaling; After a period of phase separation, the size of the cGAS-DNA condensates gradually increases and becomes sufficiently prominent to diffuse near the ER, Golgi and endosomes; At this stage, ZDHHC18 and MARCH8 can recruit the cGAS-DNA complex with significant affinity through their IDRs, catalyzing post-translational modifications of cGAS. These modifications inhibit cGAS enzymatic activity, preventing excessive activation of the cGAS signaling pathway and maintaining innate immune homeostasis.

## Discussion

cGAS is a central player in the innate immune system, acting as a DNA sensor that triggers downstream signaling pathways for antitumor or antivirus response. Classically, cGAS is known to locate in the cytoplasm and nucleus, with occasional localization on the plasma membrane, where its activity is tightly regulated to prevent unwarranted activation by self-DNA. However, our findings reveal that cGAS also localizes to the endoplasmic reticulum (ER), Golgi apparatus, and endosomes upon DNA challenge. This organelle-specific localization adds a new dimension to the regulation of cGAS activity, highlighting the complexity and precision of innate immune signaling. The spatial regulation of cGAS through biomolecular condensates has significant implications for our understanding of innate immunity. By localizing cGAS to specific organelles and sequestering it within condensates, cells can finely tune the activity of this critical sensor to ensure a balanced immune response.

This mechanism prevents the unwarranted activation of cGAS by self-DNA, thereby avoiding autoimmune responses while ensuring a robust defense against pathogens.

The enzymes ZDHHC18 and MARCH8 were previously reported to mediate post-translational modifications such as palmitoylation and ubiquitination. Our data show that these enzymes utilize their IDRs to facilitate the binding of cGAS to the Golgi and endosomes, respectively. These IDRs undergo liquid-liquid phase separation, forming biomolecular condensates that recruit cGAS and double-stranded DNA. The broader implications of our findings suggest that phase separation could be a common strategy employed by cells to organize and regulate various signaling pathways. Similar mechanisms might operate in other contexts, where the formation of biomolecular condensates helps organize cellular processes and enhance the efficiency of biochemical reactions. For example, we hypothesize that there exist many enzymes localized on organelle membranes undergo phase separation during the catalysis of substrate post-translational modifications, which will be identified in the future.

Phase separation is emerging as a fundamental mechanism for organizing cellular biochemistry. Proteins undergo LLPS to form dynamic membrane-less organelles, also known as biomolecular condensates, playing a critical role in the spatial and temporal regulation of numerous cellular processes. In our study, we identified the IDR of ZDHHC18 and MARCH8, two proteins on organelle membrane, forms biomolecular condensation, serving as hubs for recruiting and regulating activity of cGAS. Interestingly, we discovered that the phase separation of ZDHHC18’s IDR require phosphorylation of two serine residues. Moreover, the phosphorylation defeated mutant showed a dominant negative effect in the formation of phase separation in vitro, which should be noticed in the future. These results suggested a regulatory mechanism by which post-translational modifications can dynamically influence the formation of biomolecular condensates, and consequently, the activity of cGAS. Future research should aim to elucidate the detailed molecular mechanisms by which ZDHHC18 and MARCH8 mediate the phase separation of their IDRs and the specific post-translational modifications involved. Investigating other potential regulators of this process, such as kinases and phosphatases that modify these IDRs, will provide a more comprehensive understanding of how cGAS activity is controlled.

Overall, our study uncovers a novel mechanism for the spatial regulation of cGAS activity through the formation of biomolecular condensation on organelle membranes, facilitated by the IDR of ZDHHC18 and MARCH8. These findings highlight the importance of phase separation in organizing and regulating cellular signaling pathways and provide new insights into the modulation of innate immune responses. The dynamic and reversible nature of these condensates underscores their potential as therapeutic targets for diseases associated with dysregulated cGAS activity.

### Limitations of the study

The limitations of this study include the following: although MARCH8 and ZDHHC18 were identified as proteins that facilitate the binding of cGAS to endosomes and the Golgi, the underlying reasons for cGAS enrichment on the ER remain unclear. Additionally, due to limitations in research techniques, this study did not delve deeply into the molecular mechanisms by which MARCH8 and ZDHHC18 interact with the cGAS-DNA complex. Furthermore, the lack of an effective animal model hindered the in vivo investigation of the biological significance of this study.

## Materials and Methods

### Cell culture

HEK293T (ATCC CRL-3216), HeLa (NICR) and BJ-5ta (a gift from Conggang Zhang’s lab) cells were cultured in Gibco™ Dulbecco’s modified Eagle’s medium (DMEM) supplemented with 10% (v/v) fetal bovine serum (FBS), 2 mM L-glutamine, 100 U/mL penicillin and 100 mg/mL streptomycin. THP-1 lucia ISG cells (InvivoGen, thpl-isg) were cultured in Gibco™ RPMI 1640 medium supplemented with 10% (v/v) FBS, 2 mM L-glutamine, 100 U/mL penicillin and 100 mg/mL streptomycin. All cultured cells were grown at 37 °C in a humidified incubator containing 5% CO_2_. All cell lines tested negative for mycoplasma contamination.

### Reagents, plasmids and antibodies

Golgi isolation kit was purchased from Invent biotechnologies (GO-037); endosome isolation kit was purchased from Invent biotechnologies (ED-028). HT-DNA was purchased from Sigma (D6898); 17-ODYA was purchased from Cayman (90270); cGAMP was purchased from Sigma (SML1299); Anti-Flag affinity gel was purchased from Bimake (B23102). The biotin-or Cy5-labeled ISD (TACAGATCTACTAGTGATCTATGACTGATCTGTACATGATCTACA) was generated by annealing two complementary single-stranded DNA oligonucleotides.

The plasmids used in this work are listed in the key resources table. All expression plasmids were constructed by subcloning into existing vectors through homologous recombination with the pEASY-Uni Seamless Cloning and Assembly Kit (Transgen, # CU101). Point mutations were introduced with the Fast Mutagenesis System. Complementary DNAs (cDNAs) were obtained by standard PCR techniques. All constructs were confirmed by sequencing. The following antibodies were purchased from the indicated suppliers. Anti-GAPDH (CST, #2118S); anti-cGAS (CST, #15102S); anti-EEA1 (proteintech, #68065-1-lg); anti-GM130 (proteintech, #66662-1-Ig); anti-CALNEXIN (proteintech, #66903-1-Ig); TGN46 (proteintech, #13573-1-Ig); anti-Flag (MBL, #PM020); anti-HA (MBL, #PM561); Alexa Fluor 488 goat anti-mouse IgG (H+L) (Invitrogen, A11001); Alexa Fluor 488 goat anti-rabbit IgG (H+L) (Invitrogen, A11034); Alexa Fluor 568 goat anti-mouse IgG (H+L) (Invitrogen, A11031); Alexa Fluor 568 goat anti-rabbit IgG (H+L) (Invitrogen, A11011).

### OptiPrep density gradient centrifugation

To investigate the subcellular distribution of cGAS under different stimulation and lysis conditions, we performed OptiPrep density gradient centrifugation using THP-1 cells. Cells were cultured in RPMI 1640 medium supplemented with 10% fetal bovine serum and 1% penicillin-streptomycin at 37°C with 5% CO₂. For the gradient analysis, approximately 1×10^7^ cells were used per condition. Four experimental groups were prepared: (1) untreated (Mock), (2) cells transfected with 2 µg/mL HT-DNA (Sigma, D6898) for 2 hours using Lipofectamine 2000 (Thermo Fisher), (3) Mock-treated cells lysed in the presence of 1% Triton X-100, and (4) HT-DNA-transfected cells lysed in the presence of 1% Triton X-100. Cells were harvested by centrifugation at 500 × g for 5 minutes at 4°C and washed twice with ice-cold PBS. Pellets were resuspended in 800 µL of ice-cold lysis buffer containing 20 mM HEPES (pH 7.4), 0.2mM MgCl_2_, 1 mM EDTA, and protease inhibitor cocktail. The cell suspensions were incubated on ice for 30 minutes, followed by mechanical disruption by passing through a 25G needle 30 times. Lysates were centrifuged at 1,000 × g for 10 minutes at 4°C to remove cell debris.

Supernatants were further centrifugated at 5,000 × g for 20 minutes to remove nucleus and plasma membrane structure. For the relevant groups, 1% (v/v) Triton X-100 was included in the lysis buffer. The supernatants were carefully layered on top of a continuous 10%–40% OptiPrep (6%-24% iodixanol) gradient. The gradient was prepared by layering 6.5ml 40% OptiPrep solution (prepared from OptiPrep stock, Sigma-Aldrich D1556, diluted with PBS buffer, pH 7.4) into the bottom of 14 mL ultracentrifuge tubes. 6.5ml 10% OptiPrep solution was carefully added to the top of the bottom layer. The gradient was further prepared by rotated the tube by the Gradient108 (BIOCOMP) with appropriated program. The samples were centrifuged at 160,000 × g for 16 hours at 4°C in a Beckman SW41Ti rotor. Following centrifugation, 24 fractions (approximately 500 µL each) were collected sequentially from the top to the bottom of the gradient. Each fraction was mixed with 6× SDS-PAGE sample buffer, heated at 95°C for 5 minutes, and analyzed by SDS-PAGE followed by Immunoblotting.

### Split-sfGFP reporter assay

The split-superfolder GFP (sfGFP) reporter assay was employed to study protein-protein interactions in living cells. Constructs encoding β-sheet 10 and β-sheet 11 parts of sfGFP fused to cGAS were stably expressed with β-sheet 1-9 via lenti-virus package. Fluorescence complementation, indicative of protein interaction, was monitored using a fluorescence microscope. The intensity and localization of the reconstituted GFP signal provided insights into the dynamics and specificity of the interactions under DNA stimulation. To monitor the contact of cGAS with intracellular organelles, the ERM protein was fused with the sfGFP_1-9_ fragment to detect cGAS–ER membrane contact; to assess cGAS–endosome association, a tandem FYVE domain construct was fused with sfGFP_1-9_; for cGAS–Golgi contact, the N-terminal 100 amino acids of GM130 were fused with sfGFP_1-9_. As a positive control, TRAF6 was fused with sfGFP_1-9_. The negative control consisted of the human serum albumin signal peptide (MKWVTFISLLFLFSSAYSRGVFRR) fused to the N-terminus of sfGFP_1-9_ and a KDEL sequence fused to the C-terminus, ensuring retention in the ER lumen.

### Immunofluorescence

Cells were seeded on coverslips in 12-well plates and transfected with the indicated plasmids using Lipofectamine 3000 as indicated. After removing the supernatants, the cells were washed with PBS, fixed with 4% paraformaldehyde, permeabilized with 0.2% (v/v) Triton X-100 and blocked in 3% (w/v) BSA for 1 h. Then, the cells were incubated with the indicated primary antibodies (in 3% (w/v) BSA) overnight at 4 °C followed by incubation with fluorescent secondary antibodies and DAPI for 1 h at room temperature. Finally, the coverslips were fixed on slides using aqueous mounting medium, and images were acquired with a Nikon A1RMP or Zeiss LSM710 confocal microscope. Colocalization analysis was performed using Imaris software.

### Correlative Light and Electron Microscopy (CLEM)

To investigate the subcellular membrane association of GFP-cGAS, correlative light and electron microscopy (CLEM) was performed using a resin-embedding and cryo-sectioning workflow.

GFP-cGAS stably expressing HeLa cells were seeded onto poly-D-lysine– coated gridded glass-bottom dishes (MatTek, D35G-14-1.5Gl). 2-hours post transfection with HT-DNA (2 μg/ml), cells were fixed with 4% v/v paraformaldehyde (Electron Microscopy Sciences) solution in PB buffer for 30 minutes at room temperature, followed by three washes with PB buffer.

Fluorescence imaging was performed using a Zeiss LSM980 confocal microscope with a 63× oil immersion objective. The positions of cells exhibiting prominent GFP-cGAS signals were recorded based on the gridded dish coordinates.

After imaging, the samples were osmicated with 1% osmium tetroxide/1.5% potassium ferricyanidein in distilled water for 30 min. Then, samples were washed three times with distilled water and then dehydrated through a graded ethanol series. After dehydration, samples were infiltrated with Pon 812 Resin (SPI) by incubating the samples in a diluted series of ethanol-Pon 812 at a 1:1, 1:2 and 1:3 ratio for 1 h each, and then overnight in pure resin. After that, pure resin was changed once in the first 1h and then incubated in an oven at 60 °C for 48 h. 70 nm sections were cut by ultramicrotome (Leica EM UC7) then stained by uranyl acetate (UA) and lead citrate. Then cells were imaged by TEM.

Transmission electron microscopy was conducted using a FEI Tecnai Spirit TEM D1297 operating at 100 kV. Electron micrographs of regions corresponding to the previously imaged fluorescent cells were acquired at appropriate magnifications.

Correlative image processing and alignment were carried out using ImageJ software. The EM images were registered to the corresponding fluorescence micrographs by aligning plasma membrane and cGAS puncta. EM images were overlaid onto the fluorescence images to visualize the spatial relationship between GFP-cGAS puncta and organelle membranes.

### Opto-droplets activation

Cells were plated at around 70% confluency in DMEM. Constructs containing the light-sensitive protein Cry2 fused to the IDR of ZDHHC18 or MARCH8 were transfected into cells. 24 hours post transfection of opto-MEMCA^IDR^ plasmids, cells were exposed to blue light using 3min of light-dark cycles (4 s light followed by 10 s dark). The kinetics of condensate formation and dissolution were monitored in real-time using live-cell fluorescence microscopy. Imaris was used to quantify mCherry foci. Values were represented via violin plots elaborated in GraphPad Prism 8. A non-parametric t test (MannWithney) was used to compare mCherry spot/cell distributions between samples. Three biological replicates were performed in each experiment.

### TIRF-SIM

Total Internal Reflection Fluorescence Structured Illumination Microscopy (TIRF-SIM) was used to achieve super-resolution imaging of biomolecular condensates. Cells expressing fluorescently tagged proteins were plated on glass-bottom dishes. TIRF-SIM imaging was performed using a custom-built setup, allowing visualization of the nanoscale organization of condensates near the plasma membrane. High-resolution images were analyzed to determine the structural properties and spatial distribution of the condensates.

### LOV-turbo proximity labelling

For proximity-dependent labeling of proteins interacting with the intrinsically disordered regions (IDRs) of ZDHHC18 and MARCH8 condensates, we performed LOV-Turbo-based proximity biotinylation in THP-1 cells (Lee et al., 2023). Coding sequences corresponding to the IDRs of human ZDHHC18 and MARCH8 were cloned in-frame into pTy expression vector encoding a fusion protein of LOV-Turbo, Cry2, and mCherry (LOV-Turbo-Cry2-mCherry-ZDHHC18^IDR^ and LOV-Turbo-Cry2-mCherry-MARCH8^IDR^). Constructs were verified by Sanger sequencing. Subsequently, stable THP-1 cell lines expressing the above-mentioned components were generated via lentiviral packaging and transduction. Cells were cultured in RPMI 1640 medium supplemented with 10% FBS and 1% penicillin-streptomycin at 37°C and 5% CO₂.

Approximately 2×10^7^ cells per condition were collected 24 hours post-transfection. To initiate proximity labeling, cells were resuspended in RPMI medium supplemented with 500 µM biotin (Sigma-Aldrich) and exposed to continuous blue light (488 nm) for 30 minutes at room temperature using a custom LED illumination device. Immediately following light exposure, cells were transferred to ice to halt further labeling reactions. As a negative control, cells were absent from blue light exposure.

Cells were lysed in RIPA buffer (50 mM Tris-HCl pH 7.4, 150 mM NaCl, 1% NP-40, 0.5% sodium deoxycholate, 0.1% SDS, 1× protease inhibitor cocktail) for 30 minutes on ice, followed by centrifugation at 14,000 × g for 15 minutes at 4°C to remove debris. The clarified lysates were incubated with pre-washed streptavidin beads overnight at 4°C with gentle rotation. The beads were washed five times with RIPA buffer and twice with PBS to remove nonspecific binders. Bound proteins were eluted in SDS sample buffer containing 2 mM biotin at 95°C for 10 minutes.

Eluted proteins were subjected to in-gel digestion or on-bead digestion using trypsin, followed by LC-MS/MS analysis. Raw mass spectrometry data were processed with MaxQuant software against the UniProt human database, with carbamidomethylation of cysteine as a fixed modification and oxidation of methionine and N-terminal acetylation as variable modifications. Label-free quantification (LFQ) was enabled. Proteins identified in the LOV-Turbo-Cry2-mCherry-ZDHHC18^IDR^ or –MARCH8^IDR^ groups were statistically compared with those from the negative control group (blue light without biotin). LFQ values were log₂-transformed and normalized. Statistical analysis was performed using a two-tailed unpaired Student’s t-test. Proteins with log₂(fold change) ≥ 2 and p-value ≤ 0.05 were considered significantly enriched. Volcano plots were generated using GraphPad Prism 8, with log₂ (fold change) on the x-axis and –log₁₀(p-value) on the y-axis, to visualize candidate interacting proteins.

### Protein expression and purification

Recombinant proteins for in vitro assays were expressed in Escherichia coli BL21(DE3) cells. IDR of ZDHHC18 or MARCH8 and cGAS fused to His6-SUMO tags were cloned into a pET28b plasmid. The BL21 (DE3) E. coli strain harboring His6-SUMO–tagged proteins was induced with 0.5 mM isopropyl-β-D-thiogalactopyranoside (IPTG) at 18°C for 20 h.

For His6-SUMO tagged MEMCA^IDR^ expression, bacteria were sonicated in lysis buffer containing 20 mM Tris, pH 7.5, 500 mM NaCl, 25 mM imidazole, 5 mM β-mercaptoethanol, and 0.2 mM phenylmethylsulfonyl fluoride (PMSF). After centrifugation at 12, 000 rpm for 60 min, the supernatant was incubated with Ni-NTA beads (GE Healthcare), washed with lysis buffer, and eluted with 20 mM Tris, pH 7.5, 500 mM NaCl, and 250 mM imidazole. For cGAS purification, after SUMO protease (Ulp1) digestion overnight at 4°C, untagged cGAS was purified on a HiTrap heparin column (GE Healthcare) with a gradient of 0.5 to 1 M NaCl in 20 mM Tris-HCl, pH 7.5. All proteins were followed by size-exclusion chromatography using a buffer of 20 mM Tris-HCl, pH 7.5, 300mM NaCl on a Superdex200 10/300 column (GE Healthcare). Fractions were analyzed by SDS-PAGE, and relevant fractions were combined and concentrated for further use.

### In vitro phase separation assay

Phase separation of purified MEMCA^IDR^ was performed in reaction buffer (20mM Tris pH 7.5, 150mM NaCl, 5 mM MgCl_2_). Reaction was initiated by mixing purified proteins at various concentrations with crowding agents like polyethylene glycol (PEG) or ficoll by gently tapping the Eppendorf. The formation of liquid droplets was monitored using confocal microscopy.

### In Vitro FRAP Assays

In vitro FRAP assays were performed as previously described with minor adjustments (Shi et al., 2022a). FRAP experiments were performed with a Nikon A1RMP confocal microscope at room temperature. MEMCA^IDR^ spots of ∼ 1-μm diameter were photobleached with 10% laser power for 0.5 seconds using 488-nm or 561-nm lasers. Time-lapse images were acquired over a 10-min time course after bleaching at 6-second intervals. The fluorescence intensities of regions of interest (ROIs) were corrected by unbleached control regions and then normalized to the prebleached intensities of the ROIs. The corrected and normalized data were fit to the single exponential model by GraphPad Prism 7.

### Rapamycin-mediated dimerization system

To investigate how subcellular localization and biomolecular condensate formation influence cGAS activity, we employed a rapamycin-inducible heterodimerization system based on FRB-FKBP12 interactions in THP-1 cells. The fusion protein eGFP-cGAS-FRB was stably expressed with various mCherry-FKBP12 fusion constructs designed to target different membrane compartments: mCherry-FKBP12 (cytosol), mCherry-ERM-FKBP12 (plasma membrane), mCherry-FKBP12-2×FYVE (endosome), and GM130(1–100)-mCherry-FKBP12 (Golgi apparatus). The GM130(1–100) fragment encodes the N-terminal 100 amino acids of human GM130, a well-characterized Golgi-localizing sequence.

All constructs were cloned into pTY lentiviral vectors under the control of EF1α promoter. Lentiviruses were generated by co-transfecting HEK293T cells with the transfer vector and packaging plasmids (psPAX2 and pMD2.G) using Lipofectamine 2000. Supernatants were harvested at 48 and 72 hours post-transfection, filtered through 0.45 µm syringe filters, and used to transduce THP-1 cells in the presence of 8 µg/mL polybrene. Transduced cells were selected with puromycin (2 µg/mL) for 7–10 days, and stable expression of both fusion proteins was confirmed by fluorescence microscopy and immunoblotting.

For induction of condensates, stably transduced THP-1 cells (2×106) were incubated with 100 nM rapamycin (Sigma-Aldrich) for 30 minutes at 37°C to induce heterodimerization between eGFP-cGAS-FRB and the respective mCherry-FKBP12 fusion proteins. Condensate formation and relocalization of cGAS were confirmed by confocal fluorescence microscopy. Following dimerization, cells were stimulated with 2 µg/mL HT-DNA for 6 hours using Lipofectamine 2000 to activate the cGAS-STING signaling pathway. cGAS activity was indicated by cGAMP production measured by cGAMP bioassay.

### cGAMP bioassay

To evaluate cGAS enzymatic activity, a cGAMP bioassay was performed. After stimulation, cells were lysed in hypotonic lysis buffer (10 mM Tris-HCl pH 7.5, 10 mM NaCl, 1.5 mM MgCl₂, 1× protease inhibitor cocktail) by repeated freeze-thaw cycles. The supernatant was boiled at 95°C for 10 min and centrifuged at 12,000g for 10 min. The supernatant was added in THP1-lucia ISG cells pre-seeded in 96-well plates together with PFO protein (1:20000, home-made). 18 hours later, secreted luciferase activity was then measured using the Quanti-Luc detection reagent (Invivogen) according to the manufacturer’s protocol. Luminescence intensity was quantified with a microplate reader and used as a surrogate readout for intracellular cGAMP levels and cGAS activation.

### Ubiquitination Assays

Ubiquitination of cGAS was performed as previously described with minor adjustments (Yang et al., 2022). Cells were seeded in six-well plates, transfected with the appropriate plasmids, and treated with MG132 (20 μM) and chloroquine (CQ) (20 μM) 5 hours before collection. Twenty-four hours after transfection, the cells were harvested, transferred to centrifuge tubes, and lysed with buffer A [50 mM tris-HCl (pH 7.5), 150 mM NaCl, 1 mM EDTA, SDS (10 g/liter), 0.5% NP-40, and 10% glycerol] supplemented with protease inhibitor cocktail on ice for 10 min. Then, a fourfold volume of buffer B [50 mM tris-HCl (pH 7.5), 150 mM NaCl, 1 mM EDTA, 0.5% NP-40, and 10% glycerol] supplemented with protease inhibitor cocktail was added, and the lysates were sonicated for 5 min in ice water until clear. The lysates were then centrifuged at 16,200g for 10 min at 4°C. 60 μl of the supernatant was kept as the whole-cell lysate, and the rest was incubated with Flag-M2 beads at 4°C overnight. The immunoprecipitants were washed three times with buffer C [50 mM tris-HCl (pH 7.5), 500 mM NaCl, 1 mM EDTA, 0.5% NP-40, and 10% glycerol], resuspended in SDS loading buffer, and denatured by heating for 10 min. The immunoprecipitants were then analyzed by Western blotting with the appropriate antibodies (4 to 15% gels were used for SDS-PAGE).

### Palmitoylation Assays

Palmitoylation of cGAS was performed as previously described with minor adjustments (Su et al., 2023). Cells were treated with 17-ODYA (Cayman, 90270) for 20h, and then were lysed in IP lysis buffer [50 mM tris-HCl (pH 7.5), 150 mM NaCl, 0.2% Triton X-100, and 10% glycerol] supplemented with protease inhibitor cocktail and phosphatase inhibitor cocktail. Cells then were disrupted using super-sonication in ice water and cell lysates were collected by centrifugation (16,200 g, 10 min) at 4°C to remove the debris. Protein concentrations were quantified by BCA assay. Cell lysates (2 mg/mL) were used to react with 1 mM CuSO4, 100 μM TBTA ligand, 100μM azide-biotin, and 1 mM TCEP for 1 h at room temperature on a rotator. Precipitated proteins were centrifuged for 5 min at 4600g. Protein pellets were washed twice with ice-cold methanol and sonicated in PBS (1.2% SDS). Samples were heated at 95°C for 5 min and diluted to a final volume of 6 mL with PBS (0.2% SDS). An aliquot of the post-clicked lysate was retained as input and the remainder was incubated with streptavidin beads on a rotator overnight at 4°C. Samples were rotated at room temperature for 2 h to resolubilize the SDS. Beads were washed five times with 0.2% SDS/PBS and placed on a rotator for 10 min in between washes. Beads were washed with ultra-pure water for three times. At this point beads were eluted in 1X sample buffer for immunoblotting.

### Luciferase reporter assay for ISG activation in THP-1 cells

To assess the impact of ZDHHC18 and MARCH8 on cGAS-mediated innate immune activation, we utilized the THP-1 Lucia™ ISG reporter cell line (InvivoGen), which stably expresses a secreted luciferase under the control of an interferon-stimulated response element (ISRE). CRISPR-Cas9 genome editing was used to generate ZDHHC18^CRISPR^ and MARCH8^CRISPR^ knockout cell lines. Knockouts were validated by immunoblotting and sequencing.

Rescue experiments were performed by lentiviral transduction of full-length (FL), ΔIDR (IDR-deletion), or IDR-truncated constructs of ZDHHC18 and MARCH8. Transduced cells were selected using puromycin (2 μg/mL) for 7 days to ensure stable expression.

For reporter assays, 5×10^5^ THP-1 ISG cells were seeded per well in a 24-well plate. Cells were transfected with 2 μg/mL herring testis DNA (HT-DNA, Sigma-Aldrich) using Lipofectamine 2000 (Thermo Fisher) or mock treated with reagent only. After 18 hours of incubation at 37°C with 5% CO₂, supernatants were collected, and secreted luciferase activity was measured using QUANTI-Luc™ reagent (InvivoGen) in a microplate luminometer (BioTek Synergy). Relative luminescence units (RLU) were normalized to unstimulated controls and expressed as fold induction. Four independent biological replicates were performed for each condition. Statistical significance was calculated using one-way ANOVA with post hoc multiple comparisons (GraphPad Prism 8).

### Statistical Analyses

GraphPad Prism 7 software was applied to perform statistical analyses. All experiments were performed with at least triplicate samples as independent biological replicates. Two-tailed Student’s t test was used to compare data points. Significance levels are shown with asterisks, as follows: *, P<0.05; **, P<0.01; ***, P<0.001; ****, P<0.0001. Data with error bars are presented as the mean values with the SEMs.

## Acknowledgements

The authors are grateful to Professor Wei Qin for sharing the LOV-turbo constructs; to Shuaiting Yan for assistance in proximity labelling; to Xincheng Zhong for protein preparing; to Meng Han for LC-MS/MS quantification; to Jing Li and Bingyu Liu for microscopy image analysis; to Wei Liu, Tian Chen and Dr. Siqi Shen for helpful suggestions; to Dr. Jingjing Guo for helpful discussion; to Ying Li (Cryo-EM Facility of Tsinghua University, Branch of National Protein Science Facility) for resin section and Yanjie Li for technical support during TME data collection.

This work was supported by funds from the National Natural Science Foundation of China (Grant No. 22137004 and 32300593), the Natural Science Foundation of Beijing,China (Grant No. IS23107).

## Author Contributions

H.Y. conceptualized the project and supervised all the experiments; C.S. designed and performed the experiments, interpreted the data and wrote the manuscript; C.S. performed the palmitoylation assay, K.Z. performed the sucrose gradient assay.

## Data and materials availability

All data associated with this study are presented in the paper and/or SI Appendix. Materials that support the findings of this study are available from the corresponding author upon request.

